# Striving towards improved full-length single-cell RNA-sequencing

**DOI:** 10.64898/2026.03.11.709548

**Authors:** Vincent Hahaut, Rebecca Siwicki, Mariana M. Ribeiro, Alice Grison, Svitlana Malysheva, Cameron S. Cowan, Simone Picelli

**Author notes:** **Corresponding author:** S. Picelli. **Disclaimer:** Please note that this manuscript is NOT going to be sent for peer review. Given some of the limitations observed, we decided to discontinue the approach presented here and focus on novel ideas. Moreover, these observations were generated between 2022 and 2023. As technology evolves fast in the long-read sequencing field, some of our observations and pitfalls may not apply to the latest long-read technology or Isoquant pipeline. We nonetheless wanted to report our preliminary results, observations, and failed attempts, as we believe they can still benefit the research community.

## Abstract

Full-length single-cell RNA-sequencing (scRNA-seq) methods provide superior transcript coverage and isoform resolution compared to their 3’-end counterparts, but are typically limited to short-read platforms. Here, we report efforts to improve the FLASH-seq protocol and adapt it for long-read sequencing on the Oxford Nanopore Technologies (ONT) platform (FLASH-seq-ONT). We developed two plate-barcoding strategies enabling higher multiplexing: a custom PCR-ligation approach (PCR-LIG) and ONT native barcoding (NB-ONT). To support data processing, we built FSNanoporeR, a comprehensive bioinformatics pipeline for barcode demultiplexing, chimeric read detection and splitting, UMI extraction, and transcript quantification. Both barcoding strategies produced high-quality transcriptomic data from HEK293T cells, with notable differences in read length distributions. We further demonstrated that monomeric and trimeric UMIs can be reliably detected in >82% of reads, enabling accurate molecular counting at isoform resolution. However, both multiplexing approaches exhibited also critical limitations, including high chimeric read rates and index-swapping artifacts. Our results highlight both the promise and current technical hurdles of full-length single-cell long-read sequencing, and provide a practical framework for researchers considering ONT-based scRNA-seq workflows.

## Introduction

Single-cell RNA-sequencing (scRNA-seq) technologies have revolutionized genomics, enabling scientists to build comprehensive cell atlases in health and diseased tissues across all kingdoms of life. However, to achieve such a milestone, choices had to be made: either an in-depth characterization of gene expression on a restricted number of cells (e.g., SMART-seq); or the sequencing of just the end of transcripts but with a much higher throughput (e.g., 10x Genomics). Moreover, both approaches still rely on short-read sequencing, a suboptimal choice if the goal is to characterize gene isoforms (Hsu *et al*., 2022; Pan *et al*., 2022; Al’Khafaji *et al*., 2024). These limitations are at least in part addressed by recent improvements in both quality and throughput of long-read sequencing technologies. It follows that SMART-seq-derived protocols are the ideal use case for these platforms, but to date no standardized experimental and bioinformatics pipelines exist.

In this preliminary study, we initially aimed to improve the chemistry of the original (short-read) FLASH-seq protocol and later modified it to make it compatible with long-read sequencing with the Oxford Nanopore Technologies (ONT) platform (Hahaut *et al*., 2022).

## Results

We first sought to improve the original FLASH-seq protocol by benchmarking > 100 reaction conditions, including new DNA polymerases and reverse transcriptase enzymes as well as testing several lysis buffer compositions (Supplementary Data). These tests resulted in minor improvements, confirming the quality and robustness of the original FLASH-seq mix. Of note, a small reduction in the cost per cell and reaction duration could be achieved by decreasing the reverse transcriptase concentration to 1.5 U/μl per cell (Fig. 1a-b) and reducing the PCR elongation time to 5 minutes, respectively (Fig. 1c-d). Furthermore, as previously demonstrated (Hagemann-Jensen, Ziegenhain and Sandberg, 2022), the reaction volume could be further reduced to < 1 μl using mineral oil as a cover to prevent evaporation. However, the risk of losing cells during sorting appears to increase below this threshold, most likely due to FACS misalignment (which normally go unnoticed but become critical when reproducibly hitting the centre of each well becomes critical). We found that a 1.5 μl (0.3 μl lysis, 1.2 μl RT-PCR mix, 2 μl oil) provided optimal results (Fig. 1e-f). Our tests also highlighted potential avenues for optimisation under specific conditions. For instance, single-nuclei lysis was best achieved with sodium dodecyl sulfate (SDS) inactivated with Tween-20, as demonstrated by the higher percentage of intronic reads and increased isoform length (Fig. 1g-h). We also showed that DTT could be replaced with tris(2-carboxyethyl)phosphine (TCEP) (Fig. 1i), an odorless reducing agent, and also that the master mix could be prepared in advance and frozen down, thus increasing protocol flexibility (Fig. 1j). Some of these tests will require additional validation in more complex cell populations.

**Figure 1.**
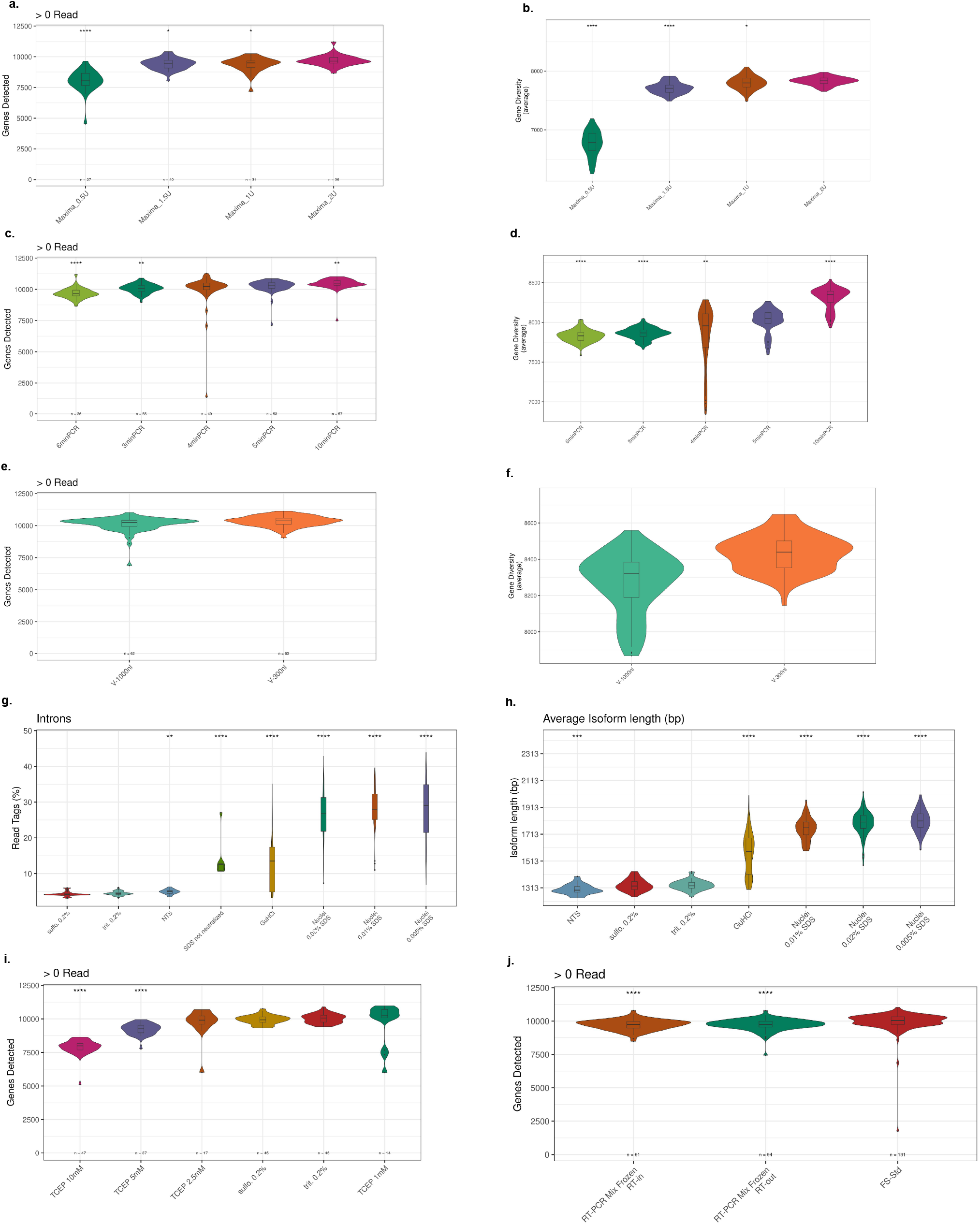
**a**. Number of genes detected for different Maxima RT concentrations. Differences in read depths were normalized by downsampling 250,000 raw reads from each cell. Only the reads mapping to the exons were taken into account. **b**. Gene diversity score for different Maxima RT concentrations. The gene diversity score calculated by selecting 10 random cells 100-times from each group and measuring the number of expressed genes in > 2 cells with > 2 reads, as in Hahaut *et al*. (2022). **c**. As in (a), but for different PCR elongation times. **d**. As in (b), but for different PCR elongation times. **e**. As in (a), but for the miniaturized RT-PCR reaction. **f**. As in (b), but for the miniaturized RT-PCR reaction. **g**. Read tag percentages for different nuclei lysis buffers. The read distribution is estimated using ReSQC. **h**. RSEM isoform length for different nuclei lysis buffers. **i**. As in (a), but for comparing different reducing agents. **j**. As in (a), but for assessing master mix viability after freezing.

We then adapted FLASH-seq to long-read sequencing on the Oxford Nanopore Technologies (ONT) platform, taking advantage of its low cost per gigabyte of data (<6$) and wide availability (Promethion P2 instrument) (Fig. 2a). We called the method FLASH-seq-ONT.

**Figure 2.**
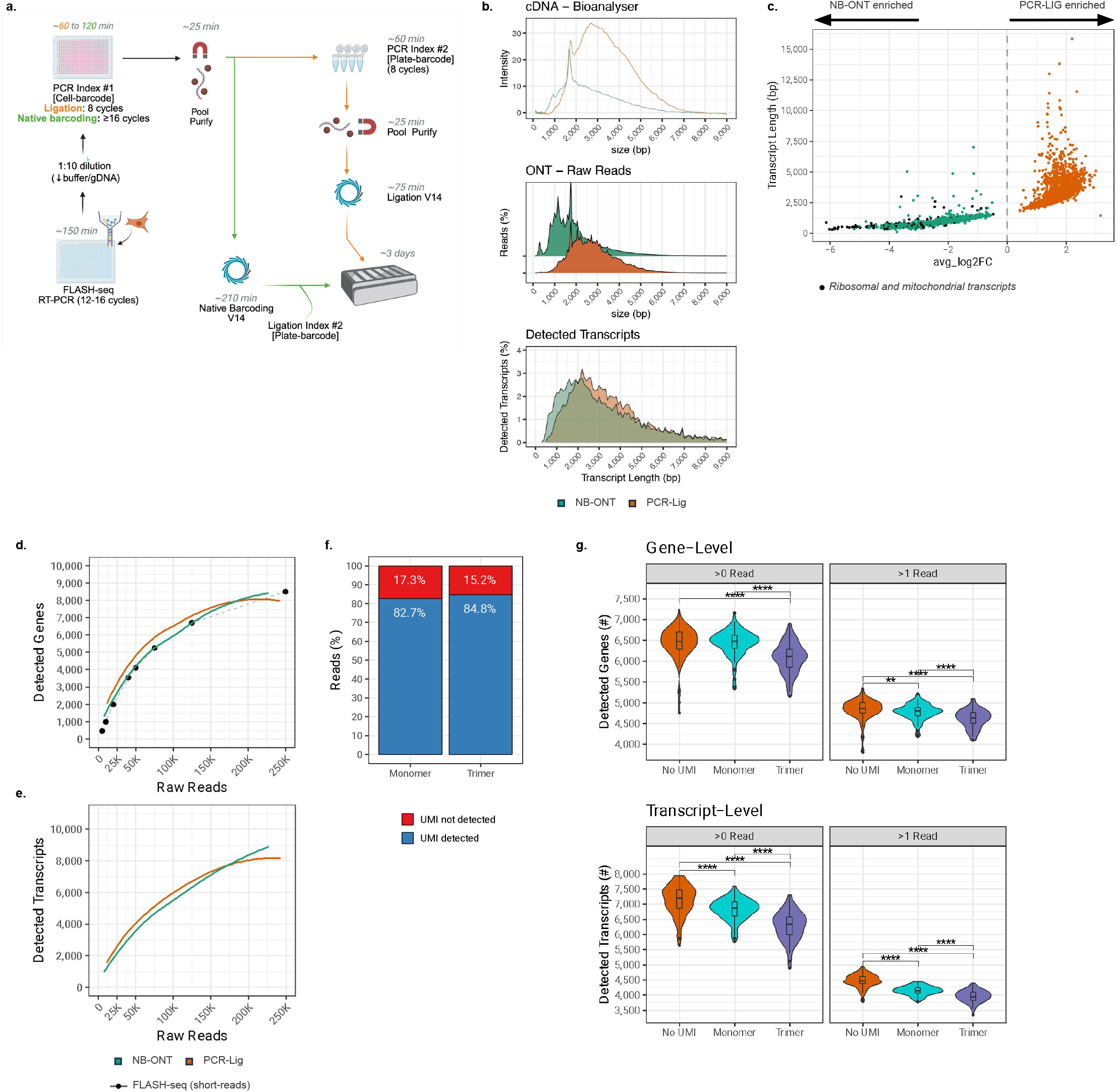
**a**. Schematic representation of the FLASH-seq-ONT protocol. After RT-PCR, the cell index is added by PCR on diluted cDNA. The number of cycle varies depending on the subsequent protocol. Indexed cDNA are then pooled and purified with magnetic beads. Multiple plates can be pooled by the addition of a plate barcode, using either ONT native barcoding kit (V14) which relies on consecutive ligations steps or a home-made PCR indexing strategy followed by ONT ligation kit (V14). Libraries are sequenced on a Promethion flow cell. **b**. Length distribution of the BC1-cDNA (Bioanalyzer, upper panel), sequenced reads (ONT, middle panel) and detected transcripts (50,000 raw reads downsampling, ONT-BC [n=113], PCR-LIG [n=93], bottom panel). **c**. Differential transcript expression between the PCR-LIG and NB-ONT cells. Average log2-fold changes as a function of the differentially expressed transcript. **d**. Number of detected genes (>1 read) with FLASH-seq-ONT as a function of the cell’s sequencing depth. A locally weighted polynomial regression was fitted to display the general trend. HEK 293T FLASH-seq short-read data from Hahaut *et al* (2022) were downsampled and the median number detected gene is displayed. **e**. Number of detected transcripts (>1 read) with FLASH-seq-ONT as a function of the cell’s sequencing depth. A locally weighted polynomial regression was fitted to display the general trend. Both unique and selected inconsistent reads were used (Isoquant). Limited to non-ribosomal and non-mitochondrial protein coding or long non-coding RNA transcripts. **f**. Percentage of reads with detected monomeric or trimer UMI. **g**. Number of detected genes and transcripts depending on the oligo-dT (50,000 raw reads downsampling, No-UMI [n=93], Monomeric UMI [n=84], Trimeric UMI [n=58]). Performed with the raw reads (no UMI). Wilcoxon’s rank sum test, two-sided, Bonferroni correction for multiple testing (adj. *P*-value, **** < 0.0001, *** < 0.001, ** < 0.01). Limited to non-ribosomal and non-mitochondrial protein coding or long non-coding RNA transcripts. Both unique and selected inconsistent reads were used for the transcript-level (Isoquant).

Following RT-PCR, the cDNA from each cell was diluted 10 times, to limit the genomic DNA carryover and dilute leftover reagents. We then designed a set of 384 error-robust 13-bp cell barcodes (BC1) to index each cell by performing a second PCR reaction. These barcodes have a Levenshtein distance of at least 4, no homotrimers, ≤ 2 homodimers and a GC content of 35-60%. FLASH-seq-ONT cell barcoding benefits from the SMART-seq semi-suppressive PCR design (Zhu *et al*., 2001), resulting in the integration of the same cell barcode at both ends, thus increasing the chance of detecting it in the analyses. After cell barcoding, cDNA from individual cells were pooled and purified with magnetic beads.

The first iteration of the protocol involved a dual barcoding system, to enable higher multiplexing capabilities. Following pooling, we then added a second 24-bp “plate” barcode (BC2). We performed an in-depth comparison of two plate-barcoding approaches on the same HEK 293T cells. We used either the ONT native barcoding ligation kit ([NB-ONT], SQK-NBD114.24) which relies on end-repair and A-tailing of the cDNA molecules, followed by ligation of the BC2. Alternatively, we carried out a PCR indexing of the BC1-cDNA molecules followed by ONT adapter ligation ([PCR-LIG], SQK-LSK114).

In parallel to this work, we developed a comprehensive pipeline to demultiplex FLASH-seq-ONT libraries (“FSnanoporeR”) as well as detect and split fusion events resulting from unwanted ligations (= chimeras) (https://github.com/vincenthahaut/FS-ONT, Fig. S1). This pipeline also integrates additional features, such as unique molecular identifier (UMI) detection and correction (monomeric or trimeric UMIs (Sun *et al*., 2024, see below), read trimming, strand assignment and optimized isoquant capabilities (Prjibelski *et al*., 2023) for gene and transcript quantifications in single cells (with or without UMIs).

Overall, both PCR-LIG and NB-ONT protocols yielded high-quality data on HEK 293T cells. The most significant difference lay in their read/transcript size distribution (Fig. 2b). NB-ONT reads were enriched in molecules < 1,000 bp in length (1,731±1,316 bp, median±SD) while the PCR-LIG samples were enriched in molecules > 2,000 bp (2,634±1,127 bp, median±SD), but both protocols were also able to capture transcripts longer than 4 kb (Fig. 2c, S2). In both cases we observed the formation of chimeric reads, derived from the combination of multiple indexed molecules (Fig. S3a-d). These appear to originate from ONT ligation(s) or are created *in-silico* when duplexes pass too quickly through the pore and are spuriously considered as one single molecule (White *et al*., 2017). Chimeric reads were more frequent in the NB-ONT protocol in this first experiment (PCR-LIG: 0.6%, NB-ONT: 10.94%). However, subsequent PCR-LIG tests also displayed large variations in the proportion of concatenated molecules (from 0 to > 25%).

So far we have not been able to determine the factor influencing these chimeric events. We hypothesize that it could arise from the number of ligation steps (for the NB-ONT variant); differences in the ligase used (T4 ligase vs blunt/TA ligase); lack of titration [sequencing-adapters]:[input cDNA] in ONT kits; our tendency to overload flowcells to maximize read output (24 pmol / run); or inaccurate estimation of the library concentration.

Given their high occurrence, we used BLAST-short to detect the position of FLASH-seq PCR primers (i.e., ISPCR). Based on the number of ISPCR sequences per molecule as well as on their orientation, we could determine whether a molecule was chimeric, and then split it into sub-segments. To validate that the splits were carried out at the correct position, the primary and secondary alignments of the chimeric reads were compared with those of the split segments (primary). The vast majority of the segments perfectly overlapped with the chimeric read alignments, validating our splits (PCR-LIG_filtered_raw_reads_: 0.297%, NB-ONT_filtered_raw_reads_: 0.93%) and allowing us to recover the barcode identity of the split reads. The cell barcode could be identified in the majority of the reads in both protocols (PCR-LIG: 92.2%, NB-ONT: 95.7%, Fig. S3e) with >80% of the reads harboring both 5’ and 3’ cell barcodes and <20% of the barcodes displayed a mismatch, allowing for an efficient demultiplexing (Fig. S3f-h) (Reads_PCR-LIG_ 67,185 ± 40,263 [n=127], reads_NB-ONT_ 102,951 ± 44,902 [n=126]). Importantly, we generally observed a limited number of reads with chimeric barcode pairs (PCR-LIG: 0.047%, NB-ONT: 0.065%), indicating that both the chimeric read split worked as expected and that no index-swapping occurred. After mapping, Isoquant was used to quantify gene and transcript expressions (Fig. 2d-e, S4a-c).

scRNA-seq presents unique challenges, such as gDNA contamination and oligo-dT/TSO artifacts, arising from the minimal amount of starting material (Hahaut *et al*., 2022). For transcript quantification, we increased the stringency of the read assignment by filtering out any read displaying major reference inconsistencies (i.e. new exons), keywords associated with gDNA contamination, and removing any read ambiguously assigned to multiple isoforms (PCR-LIG_final_assigned_reads_: 32.8%, NB-ONT_final_assigned_reads_: 31.5%) (see Methods and Fig. S5). Overall, PCR-LIG protocol detected on average more genes and transcripts than NB-ONT (Fig. 2d-e, S4a-c). The same results were observed with Bambu (Chen *et al*., 2023), another quantification software (Fig. S4d-g). Differential expression between the two protocols showed an important difference in transcript capture rate as a function of their length (Fig. 2b, S4c). Moreover, NB-ONT displayed an increased detection in mitochondrial and ribosomal transcript as well as pseudogenes (Fig. 2c, S4c).

In parallel, we focused our attention on the accurate quantification of genes and transcripts using Unique Molecular Identifiers (UMIs). According to the feedback we collected from single-cell experts, the addition of UMIs to full-length scRNA-seq protocols (Hagemann-Jensen *et al*., 2020; Hagemann-Jensen, Ziegenhain and Sandberg, 2022; Hahaut *et al*., 2022) sequenced with short-reads has proven to be challenging. The common approach relies on adding a UMI in the template-switching oligonucleotide (TSO) and performing an under-tagmentation of the cDNA, thus generating 5’ UMI-reads and internal reads without UMIs. Unfortunately, striking the right balance between UMI- and internal reads has proved to be difficult for several users (including us), as the ratio can only be established post-sequencing.

Long-read sequencing bypasses this issue by sequencing the entire cDNA molecule along with its molecular tag. Inspired by the work of Sun *et al*. (Sun *et al*., 2024), we compared the addition of either monomeric and trimeric UMIs to the FLASH-seq-ONT oligo-dT. However, we departed from the described homotrimeric UMIs for two reasons. First, runs of homopolymers are notoriously difficult to work with on the ONT platform. Second, CCC/GGG homotrimers phosphoramidites are not part of the classical catalog of nucleic acid suppliers and are therefore expensive and difficult to obtain. Therefore, we selected four trimers phosphoramidites, part of the standard codons available from most nucleic acid manufacturers (AAA, TTC, CCG, GGT), such that their Levenshtein distance is maximized as for the homotrimers.

Both monomeric and trimeric UMI sequences could be detected in >82% of the reads with our pipeline (Fig. 2f, Fig. S6), with no bias in the UMI sequences detected (Fig. S7a). Of note, we observed a great correlation between raw read and UMI counts (Fig S7b). The monomeric UMI condition showed slightly reduced efficiency compared to the no-UMI condition in detecting genes and transcripts, while the trimeric condition performed the worst (Fig. 2g, S7c-e). These differences could likely be attributed to the increased length of the trimeric oligo-dT and the lower complexity of the UMI sequence, which is likely more prone to forming secondary structures that might interfere with the RT-PCR reaction.

In summary, our results show that, when highly accurate quantification is required, trimeric UMIs can be used, similar to what was reported by Sun *et al*. (Sun *et al*., 2024). However, for most users the initial financial investment (oligo-dT_Trimer_: ~1850$, oligo-dT_monomer_: 147$) may not justify the increased accuracy and the decreased transcript detection. Moreover, only a handful of genes appeared differentially expressed between monomeric and trimeric UMI, suggesting a limited counting error (Fig S7f). For these reasons, we decided to use monomeric UMIs in all our following experiments.

Despite the apparent benefits of multiplexing, both PCR-LIG and NB-ONT protocols displayed important limitations. On the one hand, the NB-ONT kit is a sub-optimal protocol with long incubation times (more than 3.5 hours) and varying success rates in the library preparation. It also generated a lower yield and several of our test runs displayed primer short-read peaks that were not present in the cDNA. The increase in shorter molecules in the final library led to a lower number of usable reads in the final data, likely a result of the premature exhaustion of the sequencing pores.

On the other hand, the PCR-LIG approach increased the risk of index swapping, as BC1 primers could be carried over into the BC2 reaction. To prevent its occurrence, we reduced the BC1 concentration from the initial 0.8 μM (final) to 0.04 μM and performed an additional exonuclease treatment to digest any remaining primer prior to the second barcoding step (Fig. S8). Unfortunately, we later realized that this exonuclease treatment caused a shift in read length, likely due to the digestion of small molecules.

We also attempted to optimize the PCR-LIG protocol by combining both BC1 and BC2 in a single reaction, using a different set of barcoding primers (15 bp long) displaying large differences in melting temperatures (Fig. S9). Unfortunately, this approach significantly decreased the cell- and plate-barcode detection, below values that would make the protocol a viable option for long-read sequencing. In conclusion, protocol simplifications might be possible but will likely require significant optimizations.

Given the suboptimal results of these two approaches, we advise users to skip the plate barcoding altogether. In fact, FLASH-seq-ONT libraries typically generate 100-130 Gb (40-50 millions reads) per Promethion flowcells. This throughput fits perfectly the number of reads required for a full 384-well plate at our recommended read depth (~125,000 reads/cell). When needed, high multiplexing levels can be achieved with the ONT native barcoding kit.

## Discussion

In this preliminary study, we have shown that FLASH-seq can be adapted to long-read sequencing with both custom barcoding PCR and ONT native barcoding approaches, allowing for accurate molecular counting with either monomeric or trimeric UMIs while accounting for strand orientation. Our wet-lab and bioinformatics approaches are compatible with Smart-seq2 and bulk FLASH-seq (available on Protocols.io, DOI: dx.doi.org/10.17504/protocols.io.3byl4jynolo5/v2). The cost per cell is estimated at ~2 CHF on ONT (65% of which are represented by sequencing costs, assuming 50,000 reads/cell).

One of the most significant hurdles in the protocol is the amount of cDNA required for ONT sequencing, which forces the user to over-amplify the library, likely resulting in significant losses in terms of diversity. Furthermore, while providing high-quality results, the ONT technology is not as mature as short-read sequencing yet: we repeatedly faced multiple issues, ranging from low-quality flowcells and random disconnections of the sequencer, to sub-optimal library preparation guidelines. Moreover, the constant update of ONT tools such as Guppy and Dorado have significantly impacted our throughput.

However, long-read technologies expose all the issues of short-read sequencing for gene isoform reconstruction and offer an opportunity for the user to filter out protocols and sequencing artifacts. We encourage researchers to be critical about their data and to apply stringent criteria regarding gDNA contaminations, oligonucleotide invasions, and heavily truncated molecules, which could be mistakenly taken for real expression by most transcript counting tools. New read counting methods, including stringent splicing requirements and tools dedicated to scRNA-seq are urgently needed.

## Materials and Methods

### Cell preparation and sorting

HEK293T cells (CRL-3216, ATCC) were cultured in DMEM (ThermoFisher Scientific) supplemented with 10% FBS, 2 mM L-glutamine (Ambion) and 1% penicillin/streptomycin (Gibco). Cells were cryopreserved in FBS + 10% dimethylsulfoxide (DMSO). Upon thawing, 1 ml pre-warmed DMEM was added to the vial, the content was transferred to a 15-ml Falcon tube containing 10 ml pre-warmed DMEM and the cells were centrifuged for 5 min at 300 × g. The supernatant was removed, and the cells were washed twice in 1x PBS before final resuspension in 1 mL 1x PBS. Cells were strained through 70- and then 40-µm filters (pluriSelect) and stained with propidium iodide (PI, 1 mg/ml, ThermoFisher Scientific) at room temperature to label dying cells. Cells were sorted in a plate using the F.SIGHT™ OMICS (Cytena). LoBind twin.tec 384-well plates (Eppendorf) were used in all experiments.

### FLASH-seq optimization experiments

Experiments were carried out in 384-well plates using HEK 293T cells, following the original protocol, in the standard 5 μl volume, unless otherwise indicated (Hahaut *et al*., 2022). All the experimental details are described in the corresponding Supplementary File.

### FLASH-seq-ONT RT-PCR

Mineral oil (2 μl, Sigma-Aldrich) was dispensed in single wells of a 384-well plate with a Fluent 780 workstation (Tecan). After quick centrifugation (1000 x g, 10 s), 0.3 μl lysis buffer was dispensed with the I.DOT nanodispenser (Dispendix). The plate was sealed and stored at −20°C until needed. Lysis buffer was composed of 0.006 μl Triton-X100 (10%, v/v, Sigma-Aldrich), 0.072 μl dNTP mix (25 mM each, Roth), 0.0055 μl oligo-dT (variable sequence, 100 μM, IDT), 0.009 μl Recombinant RNAse inhibitor (40 U/μl, Takara), 0.0036 μl dithiothreitol (100 mM, ThermoFisher Scientific), 0.06 μl betaine (5 M, Sigma-Aldrich) and 0.1439 μl RNase-free water. After cell sorting, plates were sealed with aluminum foil seals (VWR) and immediately placed in a −80 °C freezer until ready (< 1 year). In the case of mouse retina and hPBMC, the lysis buffer did not contain oligo-dT, which was added with the I.DOT directly after thawing.

Plates were removed from the −80 °C storage, transferred to a pre-heated Mastercycler thermocycler (Eppendorf), incubated for 3 min at 72 °C and then placed on a metal block kept in an ice bucket for 5 minutes. The RT-PCR master mix (1.2 µl) was then added with the I-DOT. The RT-PCR mix had the following composition: 0.071 µl DTT (100 mM), 0.24 µl betaine (5 M), 0.014 µl magnesium chloride (1 M, Ambion), 0.029 µl Recombinant RNAse inhibitor (40 U/µl), 0.015 µl Maxima H minus RT (200 U/µl, ThermoFisher Scientific), 0.027 µl dCTP (100 mM, ThermoFisher Scientific), 0.028 µl template-switching oligonucleotide (TSO, 100 µM, IDT), 0.75 µl KAPA HiFi Hot-Start ReadyMix (2×, Roche) and 0.026 μl RNAse-free water.

The following RT-PCR program was used: 60 min at 50 °C, 98 °C for 3 min, then N cycles of (98 °C for 20 s, 67 °C for 15 s, 72 °C for 5 min). The number of cycles depended on the cell type and protocol: here N=12 was used for HEK293T cells. The exact number of cycles can be estimated as described in FLASH-seq low-amplification protocol (Hahaut *et al*., 2022 and Protocols.io, DOI: dx.doi.org/10.17504/protocols.io.yxmvmnod5g3p/v3).

### Cell Indexing (BC1)

Following RT-PCR, the cDNA was diluted 1:10 with RNAse-free water using the I.DOT. The Fluent 780 workstation was used to transfer 1.5 µl diluted cDNA into a new 384-well plate. When performing Exonuclease I tests, the enzyme mix was added prior to cell barcoding. The cDNA was combined with 0.5 µl of Exo-CIP™ Rapid PCR Cleanup Kit (NEB): 0.035 µl Exo-CIP A, 0.035 µl Exo-CIP B, 0.3 µl 5x KAPA HiFi Buffer (Roche) and 0.13 µl RNAse-free water. The mix was spun down, lightly vortexed and incubated for 8 min at 37°C before inactivation for 2 min at 80°C. In the absence of exonuclease I treatment, 2 µl of diluted cDNA was directly used for the next reaction.

Cell barcoding primers were obtained from IDT and diluted to 0.4 µM or 2 µM in a 384-well plate. Using the Fluent 780 workstation, 1.25 µl of barcoding primer was added to each cell. The 0.4 and 2 µM primer dilutions were used for the PCR-LIG and NB-ONT respectively. The PCR master mix was assembled using the KAPA HiFi HotStart PCR Kit (Roche) and dispensed with the I.DOT (0.7 µl 5x KAPA HiFi Buffer, 0.15 µl dNTP [10 mM], 0.1 µl KAPA HiFi HotStart DNA Polymerase [1 U/µl] and 0.8 µl RNAse-free water). The following PCR program was used: 98 °C for 3 min, then N cycles of (98 °C for 20 s, 63 °C for 15 s, 72 °C for 5 min), where N=8 for the PCR-LIG conditions and N=16 for NB-ONT. If performed on other cell types with the NB-ONT protocol, N may need to be increased, to reach the starting input required by ONT.

After PCR amplification, the cDNAs were collected and pooled. Magnetic bead purification was performed with 0.8:1 ratio of SeraMag Beads (GE Healthcare) containing 18% w/v polyethylene glycol (molecular weight = 8,000) (Sigma-Aldrich) (Picelli, 2019). Of note, the first incubation of beads with cDNA was done for 10 minutes at RT. Two washes with 80% ethanol were performed for the PCR-LIG protocol while only one was done for the NB-ONT protocol. The cDNA was eluted in 40 µl of nuclease-free water for PCR-LIG and 20 µl for NB-ONT. The concentration and library sizes were measured on a Qubit Fluorometer (HS dsDNA assay) and 2100 Bioanalyzer System (HS DNA chip, software vB.02.10.51764, Agilent), respectively.

### Plate Barcoding - PCR-LIG (BC2)

The plate barcode was added by PCR. First, 2 µl BC1-cDNA (1 ng/µl) was incubated with a Exo-CIP™ mix (0.5 µl Exo-CIP A, 0.5 µl Exo-CIP B, 1.4 µl 5x KAPA HiFi Buffer (Roche) and 2.6 µl RNAse-free water) for 8 minutes at 37°C and inactivated for 2 minutes at 80°C. The PCR reaction was then assembled by adding 5.5 µl of KAPA HiFi HotStart PCR mix (1.1 µl 5x KAPA HiFi Buffer, 0.3 µl dNTP [10 mM], 0.25 µl KAPA HiFi HotStart DNA Polymerase [1 U/µL], 0.75 µl plate barcoding primer [10 µM, IDT] and 3.1 µl RNAse-free water). The following PCR program was used: 98 °C for 3 min, then 8 cycles of (98°C for 20 s, 63°C for 15 s, 72°C for 5 min). The concentration and size of the libraries were measured on a Qubit Fluorometer (dsDNA assay) and 2100 Bioanalyzer System (HS DNA chip, Agilent), respectively.

To reach the required input for ONT (100-200 fmol) we split the reaction in 8 separate tubes rather than increasing the number of PCR cycles. To decrease processing time, multiple plates were processed in parallel, each on a separate column of a 96-well plate.

The Ligation Sequencing Kit V14 (SQK-LSK114, ONT) was used to prepare the final library. When multiplexing samples, equivalent concentrations were used. We started the library preparation with >150 fmol. The 0.4:1 magnetic beads:DNA ratio recommended by ONT was replaced with 0.8:1. The Short Fragment Buffer was used. We loaded ~12-15 fmol of the library on the chip.

### Plate Barcoding - NB-ONT (BC2)

The Native Barcoding Kit 24 V14 (SQK-NBD114.24, ONT) was used for plate barcoding. We followed the ONT “Ligation sequencing amplicons” protocol, except that each 0.4:1 magnetic beads ratio was replaced with 0.8:1. Whenever possible, we started the library preparation with the highest possible input (200 fmol). The Short Fragment Buffer was used. When multiplexing samples, equivalent concentrations were utilized. We loaded ~15 fmol of the library on the chip.

### Data Acquisition

All the sequencing runs were done on a Promethion P2 sequencer (ONT, flowcells v10.4.1) controlled with MinKNOW software (≤v23.11.4), on a Dell Precision Tower 3660 computer without GPU (Intel I7-12700K, 2×16Gb RAM 4400 MHz, 4 Tb PCIe NVMe Class 40 M.2 SSD). Runs were sheltered from light using aluminum foil, even before the introduction of ONT flow cell shields.

### Basecalling, BC2-demultiplexing and IT

Basecalling was performed on a dedicated google cloud ‘a2-highgpu-1g’ linux virtual machine (12 vCPUs/85 Gb RAM, 1x A100 40GB HBM2, 2 Tb GCP Performance SSD persistent disk) using dorado (v0.3.2+d8660a3). The basecalling DNA model ‘dna_r10.4.1_e8.2_400bps_sup’ (v4.2.0) was used for all samples except for the HEK293T PCR-LIG cells, displayed in the first figure, which had to rely on the previous version (v4.1.0). Demultiplexing was done on a separate custom google cloud linux virtual machine (48 vCPUs, 208 Gb RAM, 1 Tb GCP Standard Persistent disk). Basecalled BAM files were then transformed to FASTQ using samtools *fastq* (v1.3.1). To prevent memory crashes, the data FASTQ were then split into smaller files of 250,000 reads. For both PCR-LIG and NB-ONT protocols, guppy barcoder (v6.4.8+31becc9) was used for demultiplexing plate barcode: *--fastq_out --barcode_kits SQK-NBD114-24 --min_score_adapter 30 --min_score_barcode_rear 30 --min_score_barcode_front 30 --compress_fastq*. All data subsequent analyses were performed on a dedicated google cloud linux virtual machine ‘n1-highmem-32’ (32 vCPUs/208 GB RAM, Intel Xeon® Scalable Platinum 8173M Processor, 12TB GCP Standard Persistent disk).

### FSNanoporeR pipeline

After demultiplexing the plate barcodes (BC2), BLAST-short (v2.14.0) was used to detect PCR adapter sequences (“ISPCR”, AAGCAGTGGTATCAACGCAGAGT) (*-strand plus -word_size 11 -gapopen 1 -gapextend 1 -window_size 0 -perc_identity 75*). Reads displaying >2 ISPCR sequences or any ISPCR at >200 bp of the read edges were marked as chimeric reads. The reads were then split into segments based on the ISPCR positions and orientations. To ensure a proper cut, the split segments and flagged chimeric reads were mapped using minimap2 (v2.26, *-ub --ax-splice --junc-bed*). The primary mappings of the segments were compared with the primary and supplementary mappings of their parent chimeric read. Segments displaying at least 80% genomic region overlap with at most 1 mapping of a parent read were conserved, the others were excluded from the analysis. In addition, *in-silico* chimeric or unsplit duplex reads were identified by comparing the primary mappings of the segments originating from the same parent molecules. Any segments with almost identical mapping (>80% genomic region overlap) in opposite orientations were excluded. Afterwards, BC1 sequences were extracted using vsearch (v2.22.1), searching for the PCR-anchor_BC1_ISPCR (CAGCACCTCGACGCTCTTCCGATCT NNNNNNNNNNNNN AAGCAGTGGTATCAACGCAGAGT) sequence (*--maxaccepts 5 --strand plus --wordlength 3 --minwordmatches 12 --mincols 40 -id 0*.*7*) in either the first or last 200 bp of the reads (*seqkit subseq*, v2.4.0). Barcodes with perfect match to the barcode whitelist were set aside (*BC_perfect*). PCR and sequencing errors in the other barcodes (*BC_mismatch)* were handled by calculating their Levenshtein distance to whitelist barcodes. Matches with a distance greater than or equal to the minimal distance between the whitelist barcodes were discarded. For each *BC_mismatch*, the whitelist barcode(s) with the minimal Levenshtein distance were selected. *BC_mismatch* with >3 matches were discarded. The *BC_perfect* and *BC_mismatch* lists were then rejoined. For each read, there can only be two identical barcodes at the read start / end. Reads with the same *BC_perfect* at both ends were assigned to a unique cell barcode. Reads with different *BC_perfect* were flagged as chimeric. In other reads containing at least one *BC_mismatch*, the start and end barcodes were compared. When a unique match between the start and end barcodes is found, it becomes the cell barcode (i.e. *BC_perfect ∩ BC_mismatch* == 1). This translates into the *BC_perfect* always being used when a read contains both a *BC_perfect* and a list of *BC_mismatch*. Finally, reads lacking either start or end barcodes were associated with their *BC_perfect* or *BC_mismatch* (only a unique *BC_mismatch* to the whitelist is found). All other situations (i.e. *BC_perfect ∩ BC_mismatch* > 1) were marked as ‘Undetermined’.

This operation also retrieved the position of the TSO / oligo-dT, minus a few bases, and allowed their trimming prior to demultiplexing. Whenever UMIs are present, the oligo-dT positions are inferred based on the detection of the oligo-dT-UMI sequence to trim the UMI as well (see below). The oligo-dT position, at the start or end, of the read was also used to infer the read orientation and correct it to provide a stranded protocol. Only reads with a detected oligo-dT (~80%) undergo this procedure. As Isoquant does not yet use strand information when assigning reads to genes or transcripts, this procedure did not impact downstream analysis.

The entire pipeline can be installed in a conda environment and run using a R wrapper script. Options and pipeline features are controlled with a Yaml file. The pipeline can be downloaded from Github (https://github.com/vincenthahaut/FS-ONT), but is not actively maintained or supported. An HTML report file can be generated for each demultiplexing run (as in Supplementary Data).

### Reference Genomes

Human data was mapped onto GRCh38 (Gencode v43, primary assembly).

### Unique Molecular Identifiers

Two types of UMIs were used. Either monomeric UMIs (/5Biosg/AAGCAGTGGTATCAACGCAGAGTAC NNNNNNNNATACTGACGCT_(30)_VN) or trimeric UMIs (/5Biosg/AAGCAGTGGTATCAACGCAGAGTJJJJJJJJCT_(30)_VN [J = AAA, TTC, CCG, GGT]). The trimeric oligo-dT was purchased from IDT as non-catalogue oligonucleotide, with the trimer phosphoramidites provided by Glen Research. Both were used at the standard FLASH-seq oligo-dT concentration (0.4 μM). A schematic description of the pipeline can be found in Fig. S6. The first and last 200 bp segments of each read were extracted using seqkit (v2.4.0). Vsearch was used to identify the UMI position, filtering out sequences with 85% identity and matching length <30 bp. The sequences ISPCR-N_(24)_-CT_(10)_ and ISPCR-N_(8)_-ATACTGACGCT_(3)_ were used for trimeric and monomeric UMI respectively.

Vsearch (v2.22.1) only returns two hits (reverse and forward) per query. For each segment, the UMI position was defined as the one displaying the longest match, highest identity, with <2 open gaps and no gaps >2 bp. For monomeric UMIs, the UMI was then extracted and the missing bases (=frame shift) were masked with N. For trimeric UMI, two rounds of corrections were performed. First, the UMI sequence is extracted (+/-1bp). To account for small frame-shifts in vsearch match, the sequences are split by trinucleotides over three reading frames. These trimers are compared to the supplied four trimer sequences and errors (hamming distance = 1, R *stringdist*) are corrected. The frame maximizing the number of matching trimers is selected. UMIs with 8 recovered trimers are set aside (‘perfect trimers’) to create a whitelist. UMIs with 6-7 matching trimers and ≤2 gaps are selected for the second round which aim at correcting for bigger internal frame-shifts or errors. These UMIs are compared to the perfect trimers with vsearch (≤3 mismatches). For each UMI, the best match (length and identity) is selected.

Trimers differing between the UMI and whitelist are fully masked with ‘NNN’. At most two trimers can be masked per trimeric UMI. The trimers are then compressed to monomeric sequences (AAA ⇒ A, TTC ⇒ T, GGT ⇒ G, CCG ⇒ C). In both monomeric and trimeric UMI, RT-PCR and sequencing-errors were then handled with Umi-tools.

### Mapping and read assignment

Demultiplexed reads were mapped onto the reference genome with minimap2 (v2.26-r1175) in splice-aware mode (*-ax-splice*), providing the annotation (*--junc-bed*). Despite FLASH-seq-ONT reads being strand-specific, strand orientation was not activated (*-ub*) as Isoquant did not take it into account for its quantification at the time of analysis. Whenever required, reads were downsampled using seqkit *sample*. Samtools *view* (v1.15.1) was used to select primary mappings (*-F 2308*). Isoquant (v3.2.0) was used to assign reads to features (*--count_exons --data_type nanopore --stranded none*). Of note, both ‘unique’ and ‘inconsistent’ reads were used for quantification at the gene-level while only ‘unique’ reads were considered at the transcript-level. Isoquant defines unique as “reads that are uniquely assigned to a gene and consistent with any of the gene’s isoforms” and inconsistent as “uniquely assigned reads with non-intronic inconsistencies”.

The designation of ‘unique’ and ‘inconsistent’ reads was designed for novel isoform reconstruction. We decided to refine the nomenclature by subdividing the inconsistent reads into two categories (stringent and non-stringent) based on the description of the inconsistencies compared to the references. Reads displaying key words associated with major inconsistencies (e.g. new exons) or associated with key words often seen in gDNA contaminations events were marked as non-stringent (see Fig. S5e). Reads associated with multiple transcripts were systematically discarded. This new definition allowed us to refine the count matrices and use some of the reads displaying minor inconsistencies (‘recovered inconsistent’) (*isoquant_count_matrix_inconsistent*.*R)*. Conversely, reads associated with more spurious keywords were eliminated.

UMI reads were treated as follows. Isoquant read assignments to a feature were merged with the mapped BAM files using homemade scripts (*convert_isoquant_UMItools2*.*R* and *add_tag*.*py*), adding four new SAM tags, each describing a different type of assigned feature: gene non-stringent feature (GS), gene stringent feature (RS), transcript stringent feature (IS), full-length transcript stringent feature (TS). The UMI (UB) and cell barcode (CB) tags were added as well. The stringency describes the type of read assignments considered (stringent: ‘unique’, ‘unique_minor_difference’ and ‘recovered inconsistent’, non-stringent: ‘unique’, ‘unique_minor_difference’ and ‘inconsistent’). The full-length transcripts are defined based on the SQANTI tag (‘full_splice_match’, ‘mono_exon_match’) from Isoquant and represent alignments that match all splice junctions of a transcript. These updated BAM files were then parsed with Umi-tools (v1.1.4) to deduplicate reads per gene or transcript and per cell barcode. The final results were collapsed into count matrices (*tidy_umi_tools*.*R*).

Bambu (v3.2.6) was used with default parameters.

### Gene count processing

Low quality cells were defined as having <300 expressed genes and having a percentage of mitochondria gene content >5-times the median absolute deviation. All experimental data was processed with Seurat (v4.9.9.9045). Briefly, cell counts (gene- or transcript-level) were processed with *SCTransform*, using 1000 variable features for HEK 293T cells. Cells were clustered with UMAP (*umap*.*method=uwot*, v0.1.14) using a number of PCA variables defined based on the *ElbowPlot* (8-12). Differential gene expression was performed with Seurat *FindMarkers* (*min*.*pct = 0*.*25 to 0*.*5, logfc*.*threshold = 0*.*5*). Cell clusters were defined with *FindClusters (resolution = 1*.*5)* and refined with *FindSubCluster (resolution = 0*.*5 to 1)* whenever required based on manual verification of known markers.

### Cell Barcode Design

The main set of cell barcodes (‘V1 BC1’) was designed using R *create*.*dnabarcodes* function from DNABarcodes (v1.30.0). The V1 BC1 are 13-bp long with a sequence Levenshtein distance (seqlev) ≥ 4, at most 2 homodimers (internal or with the flanking sequences) and have a GC content of 35-65%. No homotrimers were allowed (internal or with the flanking sequences). After generating the barcode list, 384 barcodes were randomly picked (10,000-times). The final V1 BC1 set was chosen maximizing the average distance between each barcode. Due to limitations with the processing power required for the first approach, the second barcode set (‘V2 BC1’) was designed by creating 100,000 15 bp random nucleotide sequences. Similarly to V1 BC1, sequences that did not fulfill the homopolymers and GC requirements were filtered out. Sets of 384 random barcodes were drawn and their Levenshtein distances were calculated (stringdist, v0.9.10). The final V2 BC1 was chosen as the one maximizing the minimal Levenshtein distance between two barcodes (minimum_distance ≥5, mean_distance 8.9±SD1.25). See *generate_barcode_list*.*R* scripts for more details.

Plate barcodes sequences were taken from ONT native barcoding sets. All primers used in this manuscript are reported in the separate Supplementary File.

### All-in-one Barcoding

HEK 293T cells (n = 384) were processed with FLASH-seq-ONT (12 PCR cycles) and diluted 1:10 in RNAse-free water. The cDNA (2 µl) was treated with the Exo-CIP Rapid PCR Cleanup Kit as previously described in two separated 384-well plates. Two different PCR mixes with different melting temperatures requirements were tested. First, KAPA HiFi HotStart PCR mix (1 µl 5x KAPA HiFi Buffer, 0.15 µl dNTP [10 mM], 0.1 µl KAPA HiFi HotStart DNA Polymerase [1 U/µl] and 0.5 µl RNAse-free water). Second, Phusion™ High-Fidelity DNA Polymerase (1 µl 5X Phusion HF Buffer, 0.1 µl dNTP [10 mM], 0.1 µl Phusion DNA Polymerase (2 U/µl) [1 U/µl] and 0.6 µl RNAse-free water). In both cases, the barcodes consisted of 1 µl of V2 cell barcoding primer (2 µM) dispensed with the Fluent 780 workstation and 0.25 µl of plate barcode (10 µM). The following PCR program was used: 98°C for 3 min, then 6 cycles of (98°C 20 s, 51°C 15 s, 72°C 5 min), and 6 cycles of (98°C 20 s, 67°C 15 s, 72°C 5 min), 72 °C for 5 min. The concentration and size of the libraries were measured respectively on a Qubit Fluorometer (dsDNA assay) and 2100 Bioanalyzer System (HS DNA chip, Agilent). Cells (n = 192) from both plates were prepared with the ONT ligation protocol (SQK-LSK 114).

## Supporting information

FLASH-seq_optimization_tests

## Code availability

Scripts used to process the data and the FSnanoporeR pipeline are available at: https://github.com/vincenthahaut/FS-ONT

## Contributions

V.H., R.S., A.G. and S.P. performed experiments. V.H. developed the bioinformatics pipeline. V.H. and M.M.R. analyzed the data. S.M. established the gating and performed the FACS sorting. C.S.C. advised on data analysis. V.H. and S.P. conceived the method, planned and supervised the work. All the authors participated in the manuscript writing.

## Acknowledgements

We thank IOB’s IT department, and especially Marc Perea, for its support during this study.

