## Supplementary_Figures for "Striving towards improved full-length single-cell RNA-sequencing": Supplementary_Figure_1.pptx.pdf

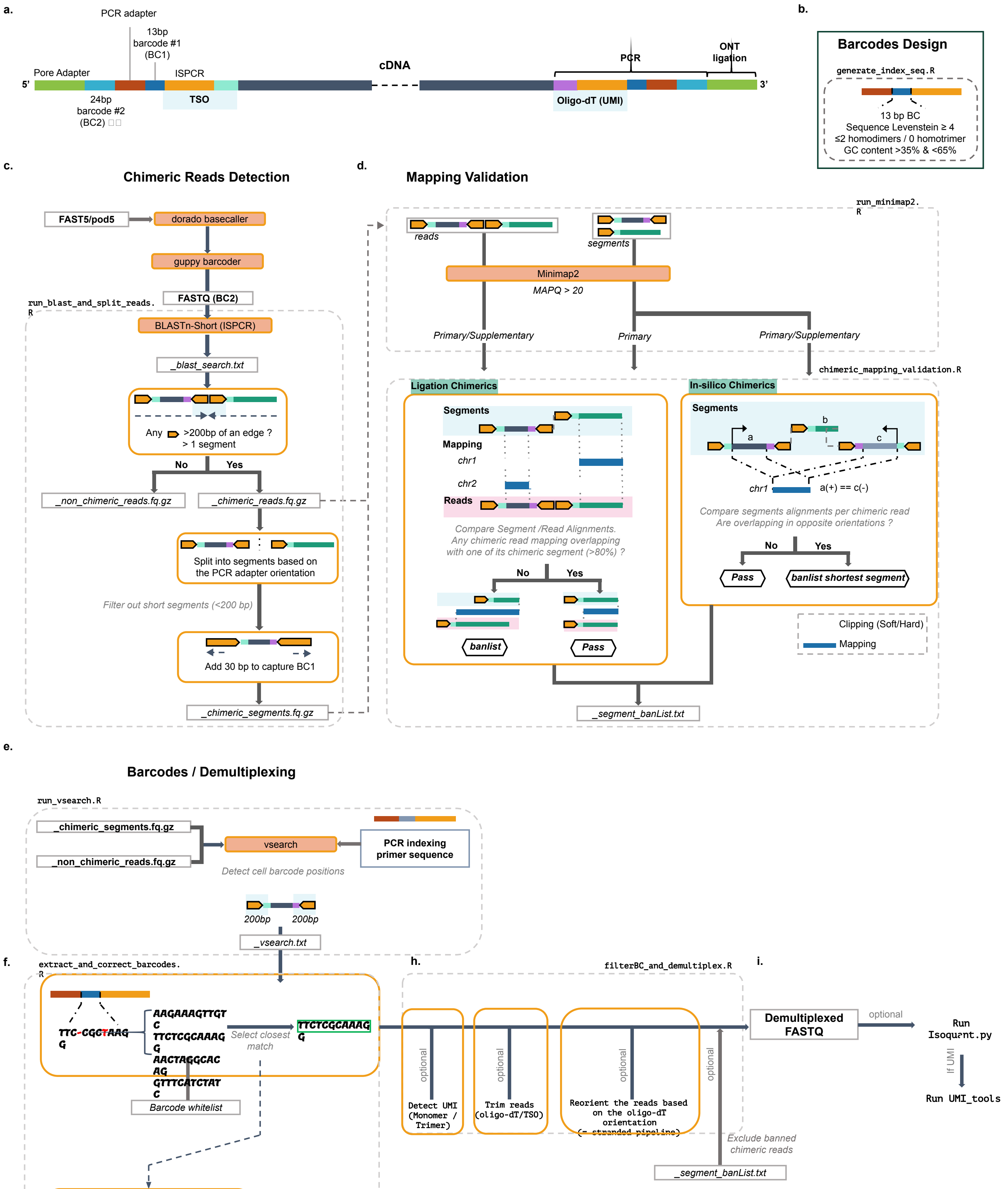


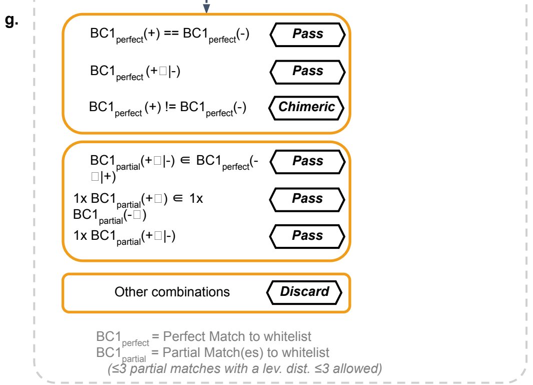


**a.** Schematic representation of FLASH-seq-ONT reads.
**b.** Criteria to design the 384-combination barcode set.
**c-i.** Schematic representation of the key aspects of FSnanoporeR pipeline.
**c.** Chimeric read detection and split, based presence and orientation of the ISPCR sequence.
**d.** Validation of the split by comparing the mappings of the split chimeric segments and associated chimeric read.
**e.** Barcode position detection using vsearch.
**f.** Barcode extraction, correction using Levenshtein distance.
**g.** Decision tree to allow mismatches in barcodes.
**h.** Demultiplexing and optional features. These include UMI detection, trimming of the oligo-dT/TSO sequences and reorientation of the reads based on the orientation of the oligo-dT.
**i.** Post-processing.
