## Supplementary_Figures for "Striving towards improved full-length single-cell RNA-sequencing": Supplementary_Figure_2.pptx.pdf

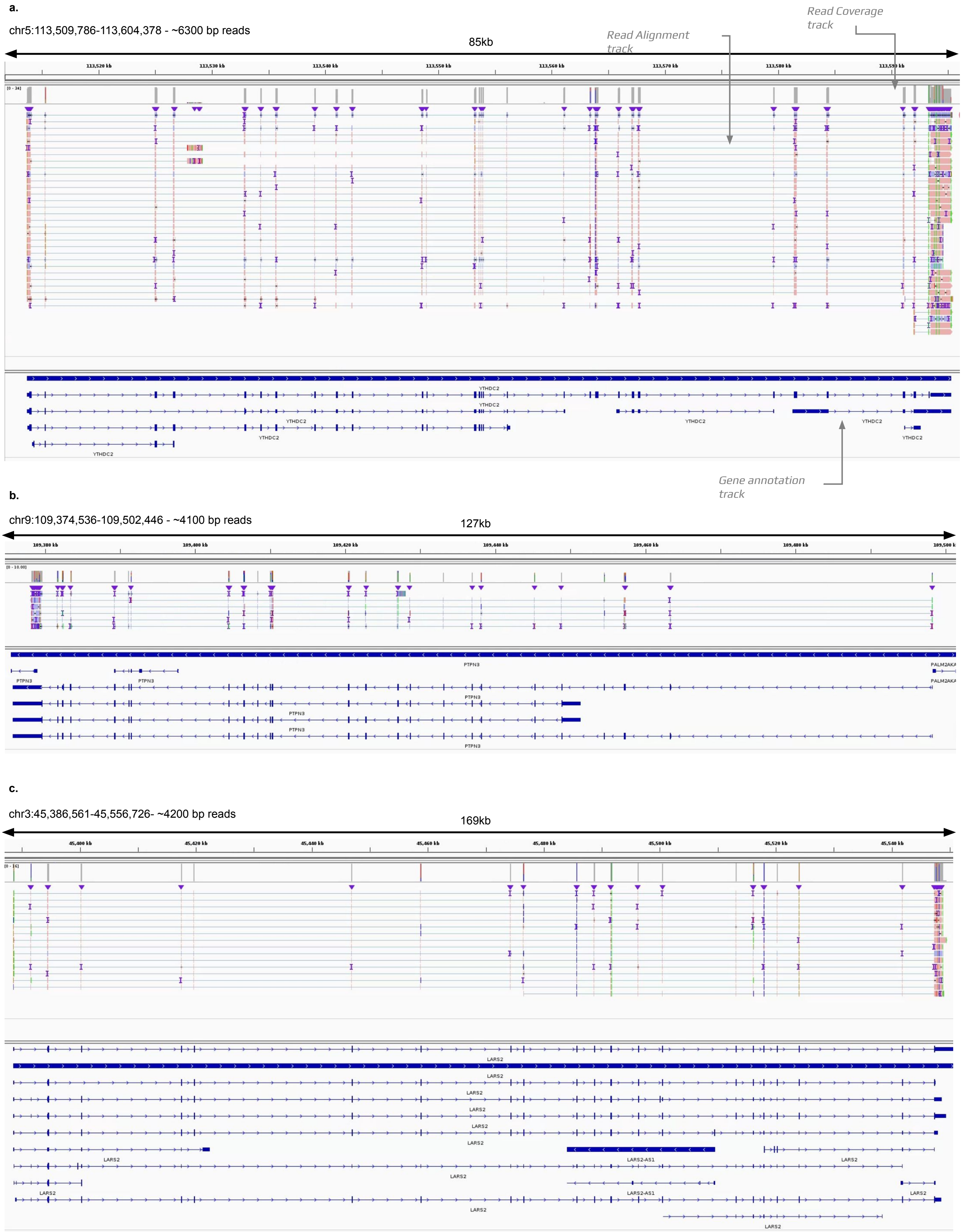


**a. – c.** Example of mapping of FLASH-seq-ONT data.
Integrated Genome Viewer (IGV) visualization of selected genes from HEK293T cells processed with FLASH-Seq-ONT (UMI – cell barcode #58).
As shown in the first panel, each sample track consists of a gene coverage and a read mapping track. In the read mapping track, each blue and red bar corresponds to a single mapped read. The color of the bar indicates the read orientation compared to the reference. Fine lines highlight split reads. The gene annotation is displayed below each panel. Fine blue lines correspond to introns and bold blue lines to exons. Each panel shows a different gene.
Some of the reads shown here are not full-length transcripts, but rather truncated fragments generated in RT-PCR, or are the result of incomplete sequencing, etc.
