## Supplementary_Figures for "Striving towards improved full-length single-cell RNA-sequencing": Supplementary_Figure_3.pptx.pdf

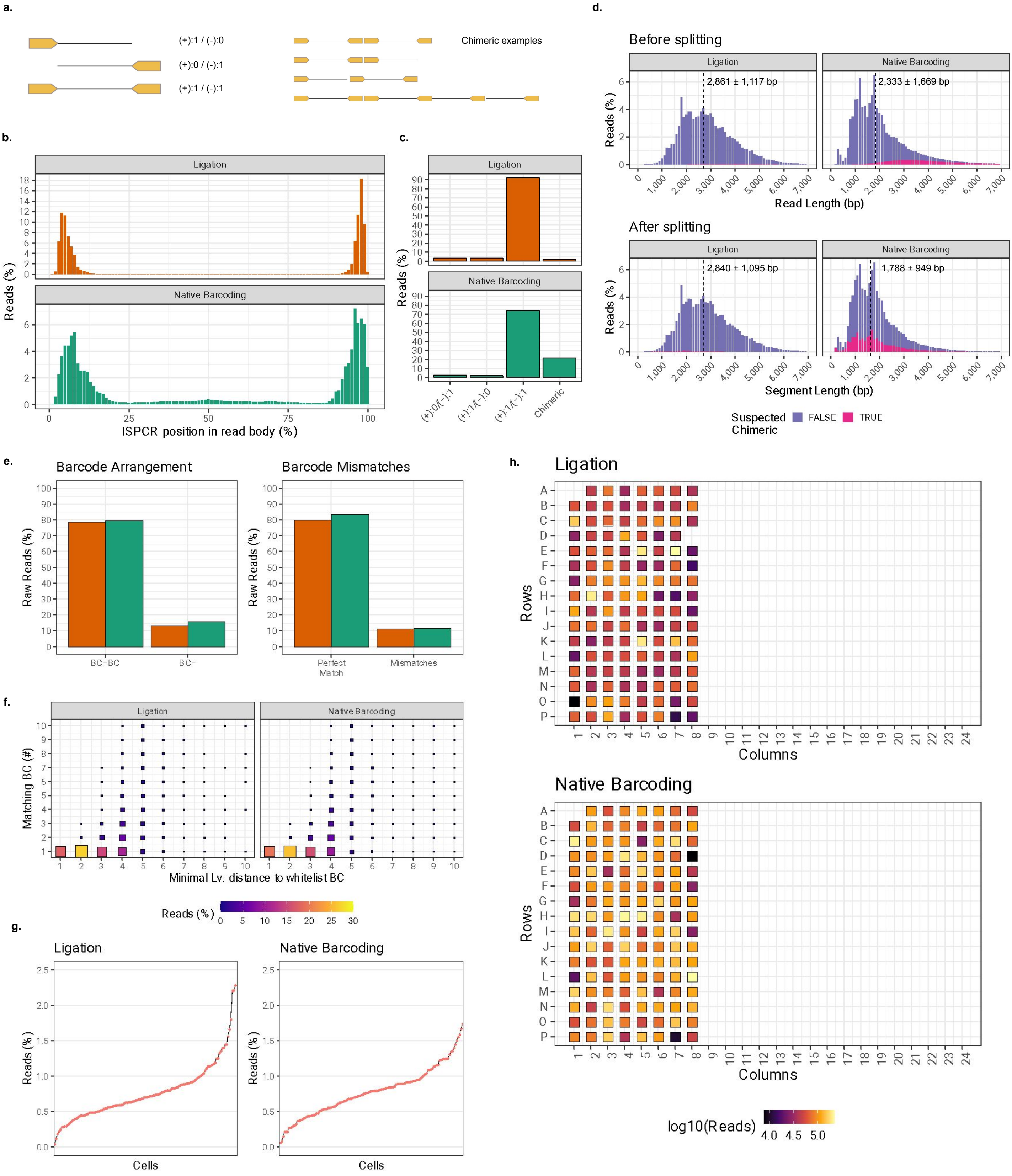


**a.** Schematic representation of the observed chimeric events.
**b.** Detected position of the ISPCR sequences inside the read body.
**c.** Number and orientation of the detected ISPCR sequences. Reads are expected to harbor an ISPCR sequence at both ends in opposite orientations (“+” and “-”). Other combinations can be considered as chimeric. Mean±SD are displayed.
**d.** Read length distribution before (top) and after (bottom) chimeric read split. Split between PCR-LIG (left) or ONT-NB (right).
**e.** Cell Barcode (BC1) detection in raw reads. From left-to right, detection of one (BC-) or two (BC-BC) barcodes per read and percentage of reads with barcodes displaying perfect matches or mismatches compared to the whitelist.
**f.** Exploration of the mismatch distribution in barcodes deviating from the whitelist. For each mismatched barcode, its levenshtein distance to the whitelist is calculated. The number of matches (y-axis) for each levenshtein distances (x-axis) is used to define if this barcode can be corrected. Only barcodes with a single match displaying a levenshtein distance of a most 3 are retained. Color and size of the square represent the percentage of barcodes with a mismatch associated to this category.
**g.** Read depth per cell post-demultiplexing.
**h.** Read depth per cell post-demultiplexing, in the context of the plate organisation. Cells were demultiplexed based on all possible barcodes (n = 384). Empty squares denote of barcodes associated with <5000 reads. Cells were sorted only in the first 8 columns of each plate.
