## Supplementary_Figures for "Striving towards improved full-length single-cell RNA-sequencing": Supplementary_Figure_4.pptx.pdf

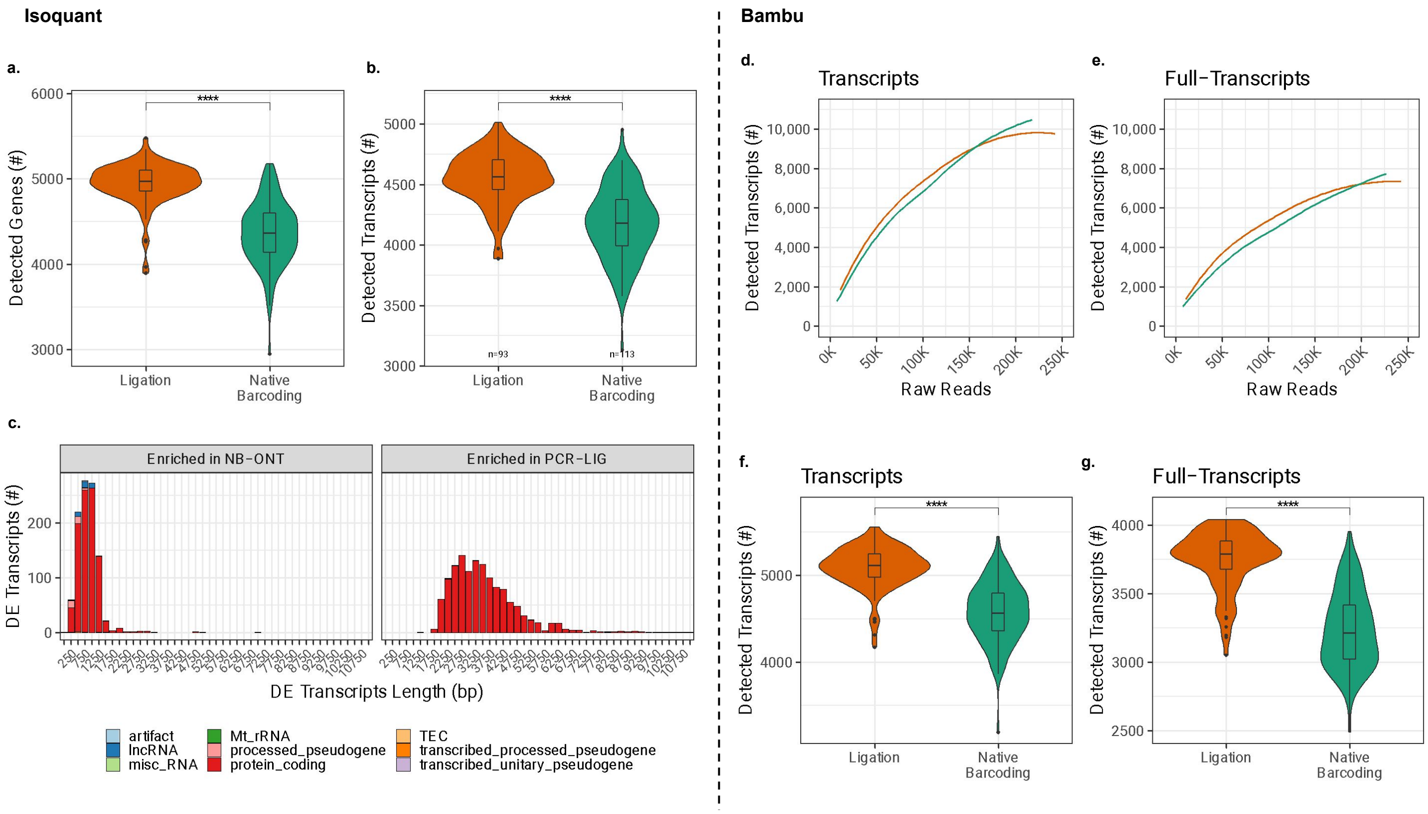


**a.** Number of detected genes (left). Cells were downsampled to 50,000 reads. Wilcoxon's rank sum test, two-sided (*P*-value, **** < 0.0001). Transcripts were assigned with both unique and selected inconsistent reads (Isoquant).

**b.** Same as (a) but for transcripts.

**c.** Transcript length distribution in both protocols. Binned by 250 bp. Gene types of the differentially expressed transcripts enriched in the NB-ONT (n = 1,023, left) and PCR-LIG (n = 1,323, right) protocols. Differentially expressed transcripts lengths were binned by 250 bp.

**d.** Transcript detection according to the cell's sequencing depth, using Bambu software.

**e.** Same as (d) but for reads mapped as "full length" for each transcript, as defined in Bambu.

**f.** Number of detected transcripts using bambu between PCR-LIG and NB-ONT conditions with cells downsampled to 50,000 raw reads. Wilcoxon's rank sum test, two-sided, Bonferroni correction for multiple testing (adj. *P*-value, **** < 0.0001).

**g.** Same as (f) but for reads mapped as "full length" for each transcript, as defined in Bambu.
