## Supplementary_Figures for "Striving towards improved full-length single-cell RNA-sequencing": Supplementary_Figure_5.pptx.pdf

a.

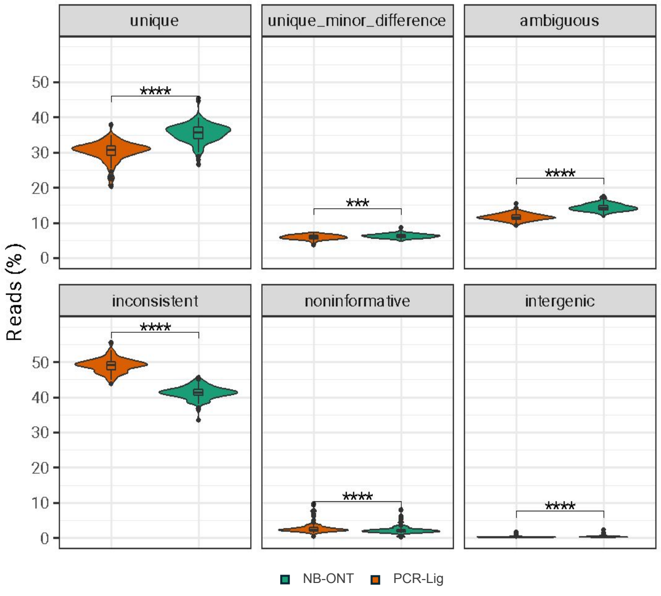


b.

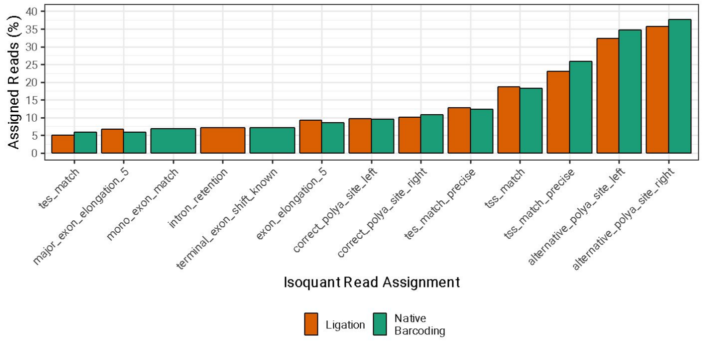


c.

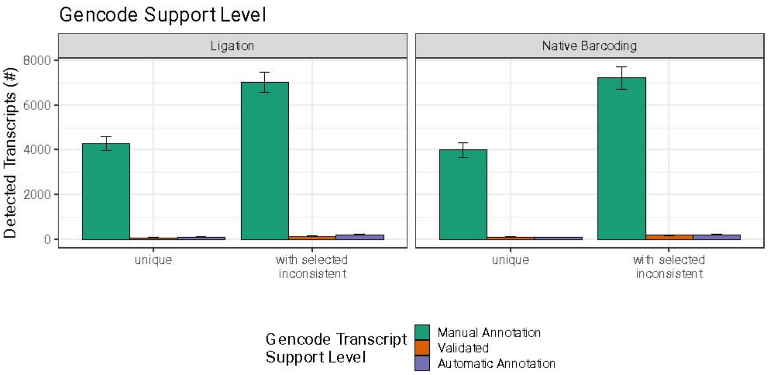


d.

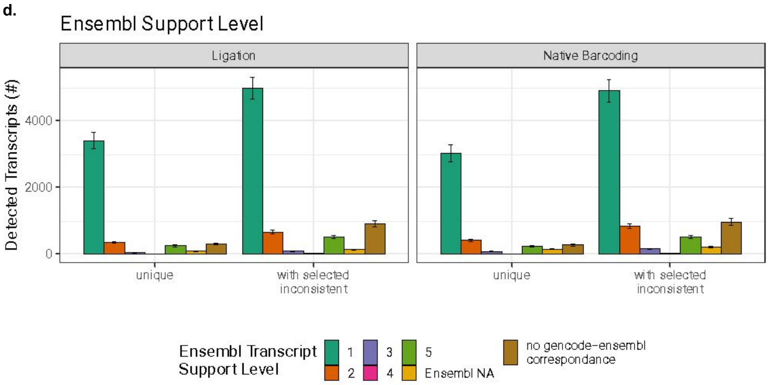


e.

| Black-listed key-words | Accepted key-words |
|---|---|
| ism (incomplete splice match) | exon_elongation |
| major_exon_elongation | alternative_polya_site_left |
| extra_intron | alternative_polya_site_right |
| alt_donor_site | exon_misalignment |
| alt_acceptor_site | terminal_exon_misalignment |
| intron_migration | fake_terminal_exon |
| intron_alternation | intron_shift |
| mutually_exclusive_exons | exon_elongation |
| incomplete_intron_retention | fsm (full splice match) |
| unspliced_intron_retention | correct_polya_site |
| intron_retention | alternative_splice_site |
| fake_micro_intron_retention | tss_match |
| exon_skipping | correct_polya_site |
| exon_merge | tes_match |
| exon_gain | aligned_polya_tail |
| exon_detach | terminal_position_match |
| terminal_exon_shift | mono_exon |
| internal_polya_site | terminal_position |
| alternative_structure | antisense |
| internal_polya | mono_exonic |
| **Classification**: antisense, genic, genic_intron incomplete_splice_match ,intergenic, novel_in_catalog ,novel_not_in_catalog | **Classification**: full_splice_match or mono_exon_match |

**a.** Distribution of the reads per transcript assignment type, as defined by isoquant.
- **unique** - reads was unambiguously assigned to a single known isoform.
- **unique_minor_difference** - read was assigned uniquely but has alignment artifacts.
- **inconsistent** - read was matched with inconsistencies, closest match(es) are reported.
- **ambiguous** - read was assigned to multiple isoforms equally well.
- **non informative** - reads is intronic/intergenic.

Wilcoxon's rank sum test, two-sided, Bonferroni adjustment for multiple testing (adj. *P*-value, *** < 0.001, **** < 0.0001)

**b.** Isoquant inconsistent features present in at least 5% of the reads. Most reveal benine inconsistent compatible with accurate read assignment.
**c.** Median (±SD) number of detected transcripts per cell, in both PCR-Lig and NB-ONT protocols. Separated by support level, according to Gencode database.
**d.** Same as (c) but for Ensembl support levels.
- **1** – all splice junctions of the transcript are supported by at least one non-suspect mRNA.
- **2** – the best supporting mRNA is flagged as suspect or the support is from multiple ESTs.
- **3** – the only support is from a single EST.
- **4** – the best supporting EST is flagged as suspect.
- **5** – no single transcript supports the model structure.
- **NA** – the transcript was not analysed for one of the following reasons: pseudogene, human leukocyte antigen transcript, immunoglobin gene transcript, T-cell receptor transcript, single-exon transcript.

**e.** Isoquant assignment events keywords. If any of the black-listed keywords are found in the description of the assigned inconsistent read, it is discarded. Otherwise, it is considered as reliable for transcript quantification.
