## Supplementary_Figures for "Striving towards improved full-length single-cell RNA-sequencing": Supplementary_Figure_6.pptx.pdf

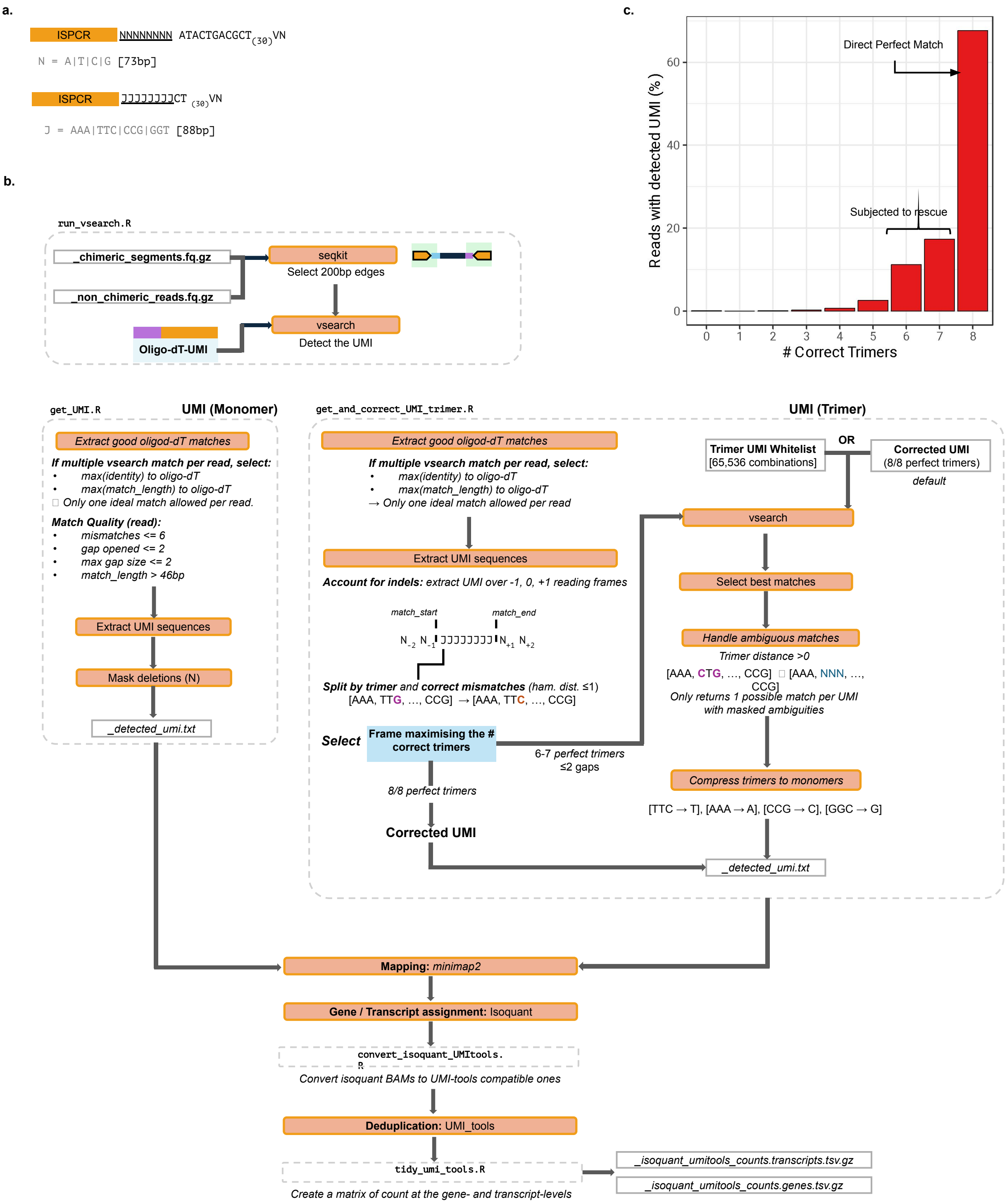


**a.** Schematic representation of the oligo-dT containing monomeric or trimeric UMI sequences.
**b.** Schematic representation of the bioinformatics pipeline to detect, extract and if necessary correct unique molecular identifiers.
**c.** Number of correct trimer per UMI as a percentage of the reads with a detected UMI.
