## Supplementary_Figures for "Striving towards improved full-length single-cell RNA-sequencing": Supplementary_Figure_7.pptx.pdf

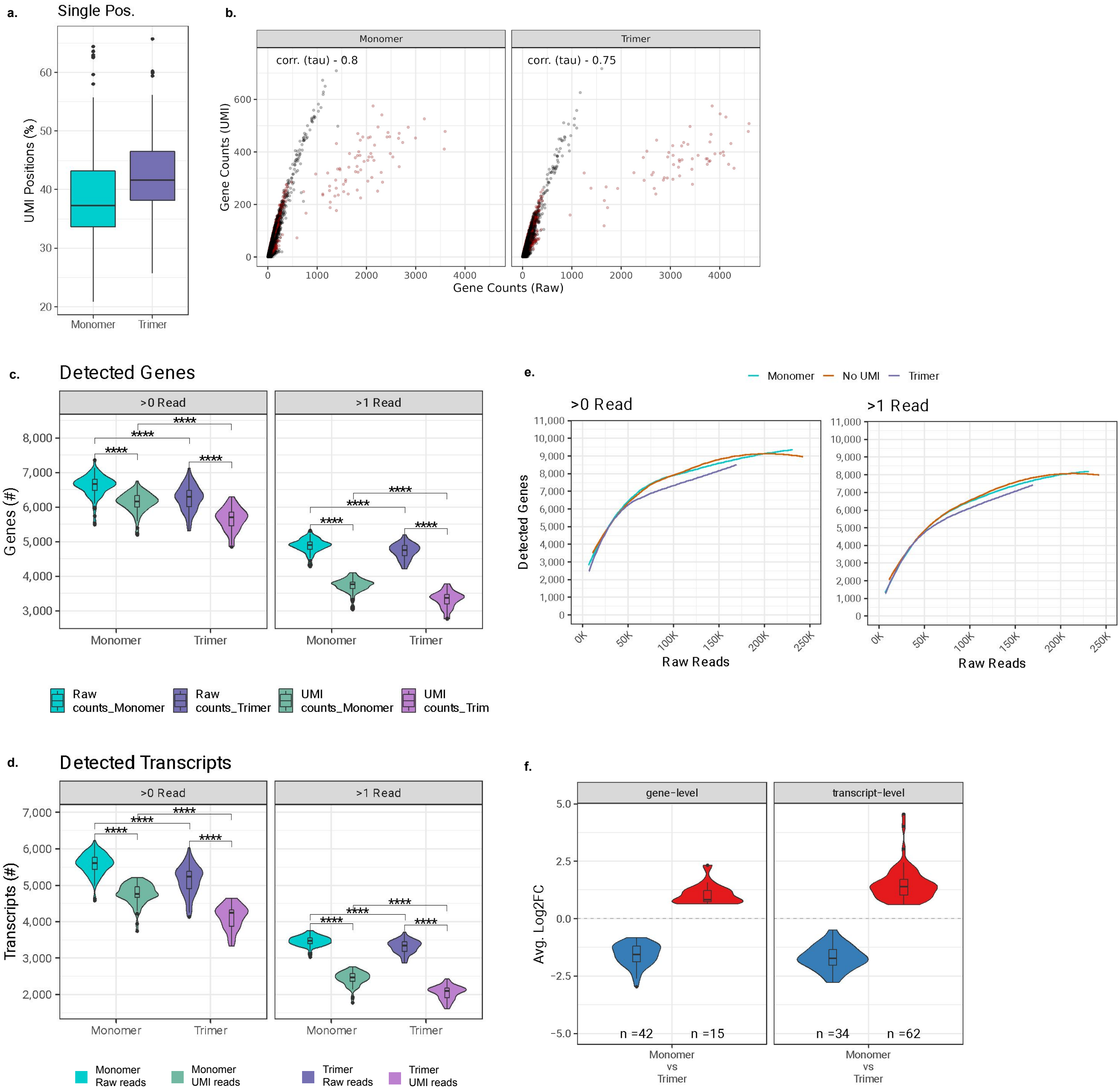


**a.** Percentage of genes associated to a single UMI sequence.
**b.** Relationship between raw read and UMI gene counts with monomeric or trimeric UMI in each cells ($n_{monomer}$=71, $n_{trimer}$=58). Cells downsampled to 50,000 raw reads. Limited to genes detected with at least one UMI. Mitochondrial and ribosomal genes are marked in red. Kendall's tau correlation displayed.
**c.** Detected genes with monomeric ($n_{monomer}$=71) or trimeric ($n_{trimer}$=58 ) UMI when accounting for all the reads (= raw reads) or UMI reads. Cells downsampled to 50,000 raw reads. Wilcoxon's rank sum test, two-sided, Bonferroni correction for multiple testing (adj. *P*-value, **** < 0.0001).
**d.** Same as (c) but for transcripts. Transcripts were assigned with both unique and selected inconsistent reads (isoquant).
**e.** Detected genes according to the sequencing depth, at two read thresholds (>0 or >1 read), without accounting for the UMI presence (= raw reads) in cells processed with monomeric UMI ($n_{monomer}$=128), trimeric UMI ($n_{Trimer}$=126) or without UMI ($n_{monomer}$=127).
**f.** Differential gene expression between the two types of UMI, performed at the gene level ($unique\_genes_{Monomer+trimer}$ = 13798) or transcript level ($unique\_transcripts_{Monomer+trimer}$ = 13798). Cells downsampled to 50,000 reads.
