## Supplementary_Figures for "Striving towards improved full-length single-cell RNA-sequencing": Supplementary_Figure_8.pptx.pdf

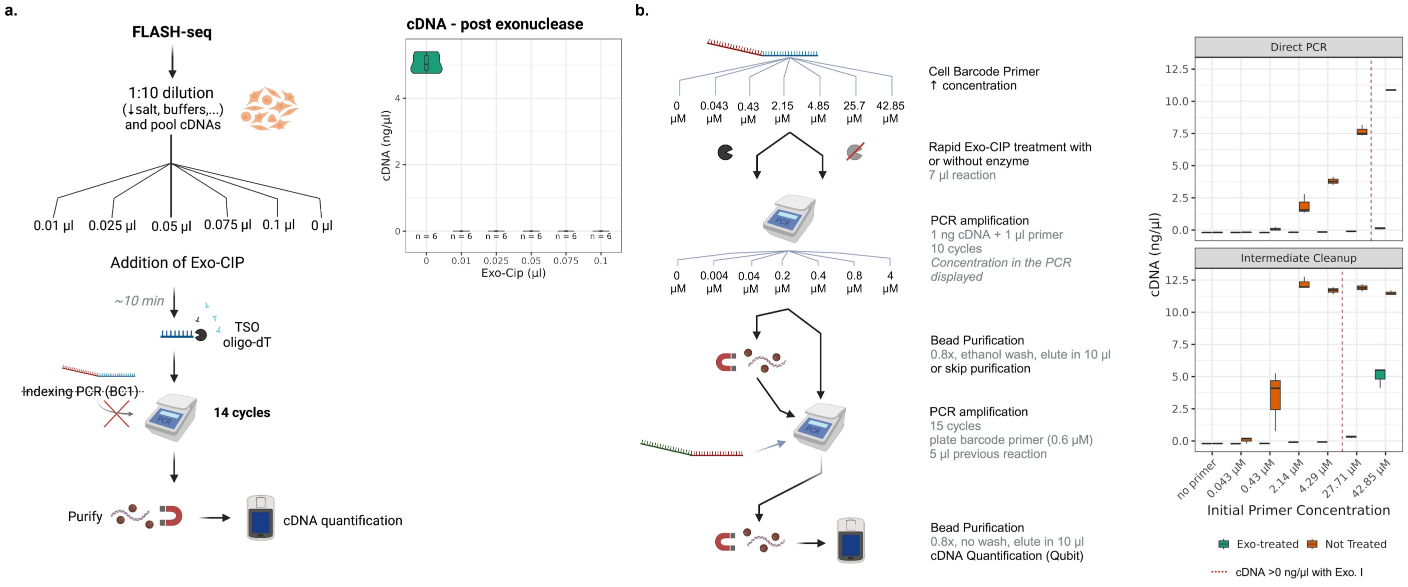


**a.** Exonuclease treatment to digest the oligo-dT and TSO after RT-PCR. cDNA from ~60 cells HEK 293T were pooled (1.84 µM TSO and 0.4 µM oligo-dT). This cDNA was treated with various amounts of rapid Exonuclease I (0 to 0.1 µl). The reaction was then incubated in the absence of additional primer for 14 cycles. After purification, the cDNA concentrations were quantified (right graphic). In the absence of treatment, the RT-PCR primer left-overs can fuel the reaction. Even the smallest tested dose of exonuclease I suppresses their use.
**b.** Rapid Exonuclease I treatment digestion of cell barcodes. Varying concentrations of cell barcoding primers (0 to 42.85 µM) were incubated w/o Rapid Exo-CIP (2 µl). As a reference, FLASH-seq-ONT cell indexing is performed with 0.1 µM primer per cell (BC1) prior to pooling and bead purification. PCR indexing was performed with 1 µl of treated primer using 1 ng of FLASH-seq-ONT cDNA (~60 cells pooled) (n = 6). These reactions were then purified with magnetic beads (‘Intermediate Cleanup’, n = 3) or left without cleanup (’direct PCR’, n = 3). This last condition was used to ensure that the purification did not result in losses of lowly represented indexed sequences. BC1 cDNA were then indexed by PCR with a plate barcode (BC2) before purification and cDNA quantification (right graphic). The red lines mark the concentrations above which the exonuclease treatment was not sufficient to prevent cDNA amplification.
