## Supplementary_Figures for "Striving towards improved full-length single-cell RNA-sequencing": Supplementary_Figure_9.pptx.pdf

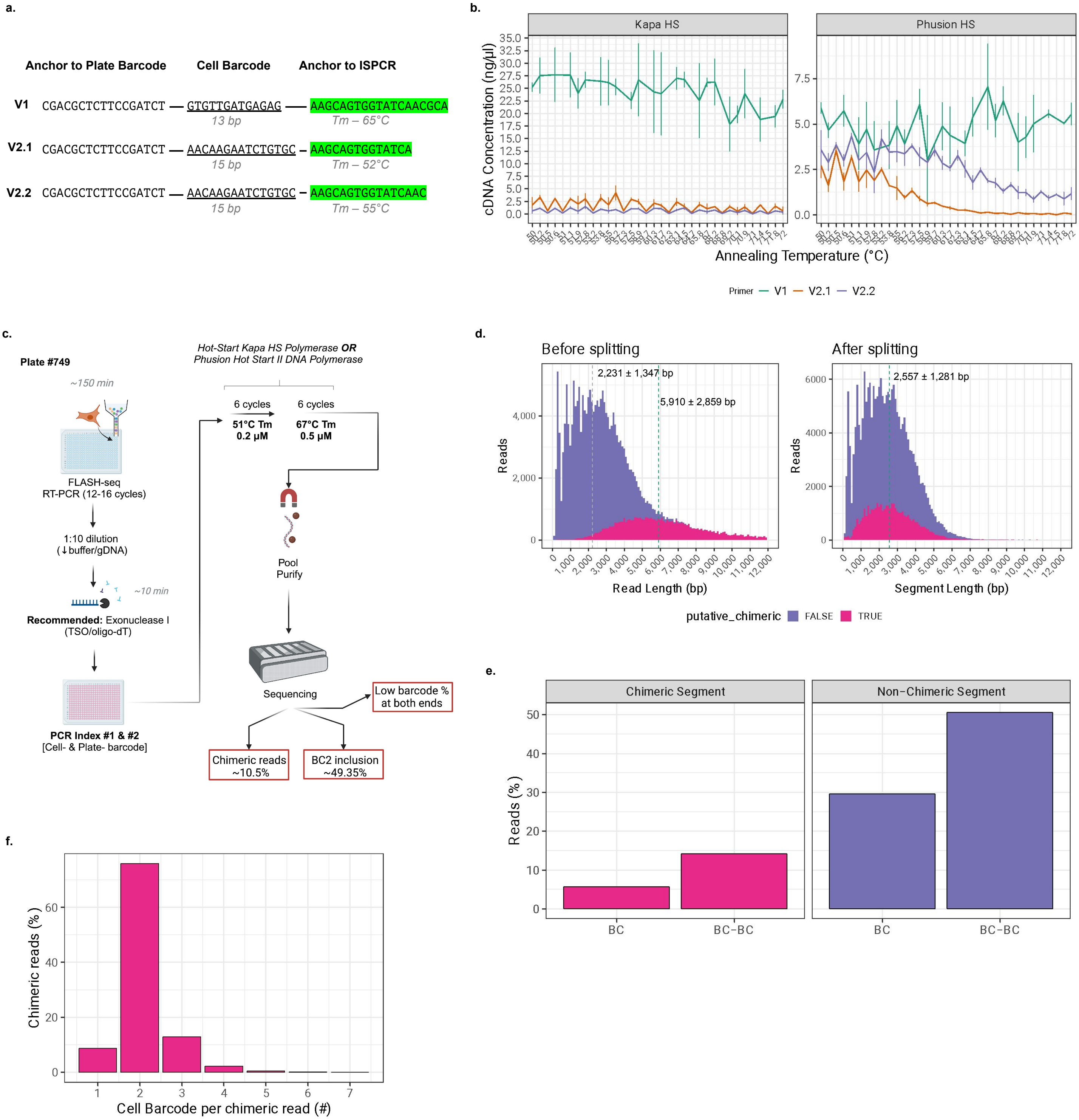


**a.** Cell indexing primer with various melting temperatures.
**b.** Gradient PCR with varying primer melting temperature from 50 to 72°C (n = 3), performed on HEK 293T FLASH-seq-ONT cDNA, using either Kapa Hot-Start Hifi polymerase or Phusion™ High-Fidelity DNA Polymerase.
**c.** Schematic representation of the all-in-one FLASH-seq-ONT protocol. Highlighting three issues: increased long chimeric reads, low inclusion of the plate barcode and low percentage of cell barcodes at both ends.
**d.** Read length distribution before (top) and after (bottom) chimeric read split. Split between PCR-LIG (left) or ONT-NB (right). Kapa Condition displayed. Median ± SD displayed for chimeric and non-chimeric reads.
**e.** Barcode detection at one end (BC) or both ends (BC-BC) of the reads in chimeric or non-chimeric reads.
**f.** Number of different cell barcode found per chimeric reads. A number greater than 1 suggests that these chimeric reads originate from the plate barcode ligation step.
