## Supplementary material for "Striving towards improved full-length single-cell RNA-sequencing": FLASH-seq_optimization_tests: DDM_as_LB.html

DDM lysis buffer - Plate 460


### DDM lysis buffer - Plate 460

#### DDM lysis buffer - Plate 460

- Introduction
- Failure
  Rate
- Raw Reads
- Mapping
  Stats
- Mapping
  Errors
- Genes
  Detected
- Isoforms Detected
- Gene
  Diversity
- C.-to-C.
  corr.
- Read
  Distribution
- Gene type
- Gene Length
  - Gene-Level
  - Isoform-Level
- Mitochondrial genes
  - Gene-level
  - Chromosome-level
- Estimated Ribosomal
  Reads
- Gene-body Coverage
- GC Content
  - Read-level
  - Gene-level
- Gene
  Expression
- Conclusions

Hahaut
V., Siwicki R., Ribeiro M., Grison A. Malysheva S. & Picelli S.

05
December 2023

### Introduction

n-Dodecyl β-D-maltoside (DDM) is a classic nonionic surfactant and
has been shown to robustly solubilize membrane proteins for effective
cell lysis. It has been shown to be an ideal buffer for single-cell
proteomics that are going to undergo Lc-MS/MS analysis. In this
experiment we replaced 0.2% Triton with 0.2% DDM and performed the
regular FLASH-seq protocol

**References** https://www.nature.com/articles/s41467-021-26514-2 https://www.nature.com/articles/s41467-021-26514-2

This file contains an overview of our tests on the DDM lysis buffer
(1 ul lysis buffer).

- **p-value < 0.05**: “\*”
- **p-value < 0.01**: “\*\*”
- **p-value < 0.001**: “\*\*\*”
- **p-value < 0.0001**: “\*\*\*\*”

Whenever possible we compared the reference group (“**72°C
3min, trit. 0.2%**”) to the other groups using a two-sided
Wilcoxon rank sum test. Only P-values inferior to 0.05 are
displayed.

For more information regarding the analysis please contact me:
vincent.hahaut[at]iob.ch.

**Sequencing:** NextSeq550 - 75bp

**Abbreviations:**

All denaturations were performed at 72°C for 3 minutes.

- **72°C 3min DDM.** : 0.2% N-Dodecyl-beta-Maltoside
  Detergent in lysis.
- **72°C 3min GuHCl.** : Guanidine Thyocyante at 250 mM
  in lysis.
- **72°C 3min trit. 0.2%** : 0.2% Triton in lysis.
- **72°C 3min sulfo. 0.2%** : 0.2% sulfobetaine in
  lysis.

### Failure Rate

Percentage of low quality cells after sequencing. Defined as:

- less than 100K raw reads
- more than 25% unmapped reads
- less than 75K uniquely mapped reads

This only represents the failure rate among cells that were chosen
for sequencing.

### Raw Reads

Number of raw reads per cell.

### Mapping Stats

From STAR mapping statistics.

### Mapping Errors

From STAR mapping statistics.

### Genes Detected

Differences in read depths were normalized by downsampling 250,000
raw reads from each cell.

Reads were mapped to the genome using STAR and the number of genes
detected was calculated using featureCounts in unstranded mode. Only the
reads mapping to the exons were taken into account.

### Isoforms Detected

RSEM.

TPM = Transcript-per-million reads.

### Gene Diversity

The gene diversity is calculated by selecting 10 random cells
100-times from each group and measuring the number of expressed genes in
> 2 cells with > 2 reads. See Hahaut et al, 2022 for detailed
information.

### C.-to-C. corr.

Cell-to-cell correlations of the gene counts are estimated using
Kendall’s tau coefficient to account for ties. In order to compare the
values between groups, the correlations are only calculated in a set of
genes expressed with at least 1 read in >2 cells from each group (n=
21602).

### Read Distribution

The read distribution is estimated using ReSQC (Hg38, Gencode v34) .
Displayed in read tag percentages.

### Gene type

The Gene type distribution is estimated using ReSQC. Only the
percentages of lncRNA and protein coding genes are displayed as they
represent the two main categories of genes detected. miRNAs, snoRNA, …
are unlikely to be captured by the FLASH-seq protocol.

### Gene Length

#### Gene-Level

Using the average length of the genes detected in the diversity
analysis.

#### Isoform-Level

Using the length of the RSEM isoforms.

### Mitochondrial genes

#### Gene-level

Estimation the percentage of mitochondrial RNA captured by FLASH-seq,
using the number of reads uniquely mapped to the mitochondrial
genes.

#### Chromosome-level

Percentage of reads uniquely mapped to the mitochondrial
chromosome.

### Estimated Ribosomal Reads

Rough estimation of the percentage of ribosomal reads, using the
reads uniquely mapping to the ribosomal genes.

As this estimation does not account for multi-mapped reads, the
percentage of captured ribosomal RNA is highly under-estimated but
provides a trend that we find useful for method development
purposes.

### Gene-body Coverage

Estimated using ReSQC.

### GC Content

#### Read-level

Average read GC content per group, measured with ReSQC (directly on
FASTQ).

#### Gene-level

Average read GC content per group. Estimation based on the average GC
content of the genes from the diversity analysis.

### Gene Expression

A first overview of the relationship between the cell’s
transcriptomes. This graph is rarely (if ever) used for defining what is
the best condition.

1. Genes supported by a total of =<10 reads among all the cells are
   discarded. “MT-|MALAT1|RPS|RPL” genes are discarded
2. Depth normalisation is performed with librarySizeFactors /
   quickCluster / computeSumFactors.
3. Counts are log-normalised (logNormCounts)
4. Top 10% highly variable genes are selected (modelGeneVar,
   getTopHVGs)
5. PCA/UMAP are run (10 PCA)

### Conclusions

DDM as a lysis buffer does not perform better than triton 0.2%.

However, the difference is often minor and could therefore replace
triton if mass spectrometry is performed downstream of FLASH-seq.
