## Supplementary material for "Striving towards improved full-length single-cell RNA-sequencing": FLASH-seq_optimization_tests: Freezing_RT-PCR_mix.html

Freezing Master Mix - Plates 588/589


### Freezing Master Mix - Plates 588/589

#### Freezing Master Mix - Plates 588/589

- Introduction
- Failure
  Rate
- Raw Reads
- Mapping
  Stats
- Mapping
  Errors
- Genes
  Detected
- Isoforms Detected
- Gene
  Diversity
- C.-to-C.
  corr.
- Read
  Distribution
- Gene type
- Gene Length
  - Gene-Level
  - Isoform-Level
- Mitochondrial genes
  - Gene-level
  - Chromosome-level
- Estimated Ribosomal
  Reads
- Gene-body Coverage
- GC Content
  - Read-level
  - Gene-level
- Gene
  Expression
- Conclusions

Hahaut
V., Siwicki R., Ribeiro M., Grison A. Malysheva S. & Picelli S.

05
December 2023

### Introduction

This file contains an overview of our tests on freezing the RT-PCR
master mix prior to processing.

The simplest single-cell RNA-seq protocol would enable the user to
sort cells in wells already containing everything for lysis, RT and PCR,
thus making the plates much more straightforward to handle. Here we
tested 2 possible variants of such an approach:

- Single cells were sorted in a mix containing everything for the
  lysis as well as the RT-PCR reaction (”RT-PCR Mix Frozen RT-in”). Upon
  thawing, we did not perform heat denaturation (72°C for 3 min) in order
  not to inactivate the reverse transcriptase, but rather proceeded
  directly with the RT reaction.
- Alternatively, single cells were sorted in a mix containing
  everything for the lysis as well as the RT-PCR reaction except for the
  reverse transcriptase (”RT-PCR Mix Frozen RT-out”). We reasoned that the
  enzyme might be damaged upon freezing in a different buffer than its own
  storage buffer. Of course, this would still require reagent addition
  before plate processing, but it would not require a complex mix.

Whenever possible we compared the reference group
(“**FS-Std**”) to the other groups using a two-sided
Wilcoxon rank sum test. Only P-values inferior to 0.05 are
displayed.

- **p-value < 0.05**: “\*”
- **p-value < 0.01**: “\*\*”
- **p-value < 0.001**: “\*\*\*”
- **p-value < 0.0001**: “\*\*\*\*”

For more information regarding the analysis please contact me:
vincent.hahaut[at]iob.ch.

**Sequencing:** NextSeq550 - 75bp

**Abbreviations:**

- **RT-PCR Mix Frozen RT-in**: Cells is lysed, RT-PCR mix
  added with RT enzyme, plate is frozen.
- **RT-PCR Mix Frozen RT-out**: Cells is lysed, RT-PCR
  mix added without the RT enzyme which was added after denaturation,
  plate is frozen.

### Conclusions

Surprisingly all the frozen mixes worked. However, as expected,
freezing the RT enzyme didn’t work as well as adding it fresh.

Despite its advantages, we decided to leave this condition aside from
the new FLASH-seq protocol. It will require more tests and longer
freezing conditions to make sure that the reaction can be kept frozen
for extended periods of time.

Additional cryopreservant may be tested in the future to improve the
conservation.
