## Supplementary material for "Striving towards improved full-length single-cell RNA-sequencing": FLASH-seq_optimization_tests: GnT-seq_like_conditions.html

G&T-seq like bead purification - Plate 605 and 677


### G&T-seq like bead purification - Plate 605 and 677

#### G&T-seq like bead purification - Plate 605 and 677

- Introduction
- Failure
  Rate
- Raw Reads
- Mapping
  Stats
- Mapping
  Errors
- Genes
  Detected
- Isoforms Detected
- Gene
  Diversity
- C.-to-C.
  corr.
- Read
  Distribution
- Gene type
- Gene Length
  - Gene-Level
  - Isoform-Level
- Mitochondrial genes
  - Gene-level
  - Chromosome-level
- Estimated Ribosomal
  Reads
- Gene-body Coverage
- GC Content
  - Read-level
  - Gene-level
- Gene
  Expression
- Conclusions

Hahaut
V., Siwicki R., Ribeiro M., Grison A. Malysheva S. & Picelli S.

05
December 2023

### Introduction

We tested 2 strategies to maximize cDNA recovery for the G&T-seq
protocol:

- We coupled a biotinylated SMART dT30VN oligonucleotide to Dynabeads™
  MyOne™ Streptavidin C1 (Invitrogen, cat. # 65001), “GT\_noBio”;
- We coupled both a biotinylated SMART dT30VN oligonucleotide and a
  biotinylated FLASH-seq TSO to Dynabeads™ MyOne™ Streptavidin C1,
  “GT\_Bio”.

With this experiment we wanted to test the hypothesis whether the RT
and PCR are more efficient when one or both oligos/primers are
immobilized on a surface rather than dispersed in solution.

References https://www.nature.com/articles/nmeth.3370

- **p-value < 0.05**: “\*”
- **p-value < 0.01**: “\*\*”
- **p-value < 0.001**: “\*\*\*”
- **p-value < 0.0001**: “\*\*\*\*”

Whenever possible we compared the reference group
(“**FS-Std**”) to the other groups using a two-sided
Wilcoxon rank sum test. Only P-values inferior to 0.05 are
displayed.

Downsampling in this document were done at 100K instead of 250K.

For more information regarding the analysis please contact me:
vincent.hahaut[at]iob.ch.

**Sequencing:** NextSeq550 - 75bp

**Abbreviations:**

- **GT\_Bio**: Both the TSO and the oligo-dT are on the
  beads.
- **GT\_noBio**: Only the oligo-dT is on the beads.

### Failure Rate

Percentage of low quality cells after sequencing. Defined as:

- less than 100K raw reads
- more than 25% unmapped reads
- less than 75K uniquely mapped reads

This only represents the failure rate among cells that were chosen
for sequencing.

### Raw Reads

Number of raw reads per cell.

### Mapping Stats

From STAR mapping statistics.

### Mapping Errors

From STAR mapping statistics.

### Genes Detected

Differences in read depths were normalized by downsampling 250,000
raw reads from each cell.

Reads were mapped to the genome using STAR and the number of genes
detected was calculated using featureCounts in unstranded mode. Only the
reads mapping to the exons were taken into account.

### Isoforms Detected

RSEM.

TPM = Transcript-per-million reads.

Downsampling was performed at 100K instead of 250K.

### Gene Diversity

The gene diversity is calculated by selecting 10 random cells
100-times from each group and measuring the number of expressed genes in
> 2 cells with > 2 reads. See Hahaut et al, 2022 for detailed
information.

### C.-to-C. corr.

Cell-to-cell correlations of the gene counts are estimated using
Kendall’s tau coefficient to account for ties. In order to compare the
values between groups, the correlations are only calculated in a set of
genes expressed with at least 1 read in >2 cells from each group (n=
17117).

### Read Distribution

The read distribution is estimated using ReSQC (Hg38, Gencode v34) .
Displayed in read tag percentages.

### Gene type

The Gene type distribution is estimated using ReSQC. Only the
percentages of lncRNA and protein coding genes are displayed as they
represent the two main categories of genes detected. miRNAs, snoRNA, …
are unlikely to be captured by the FLASH-seq protocol.

### Gene Length

#### Gene-Level

Using the average length of the genes detected in the diversity
analysis.

#### Isoform-Level

Using the length of the RSEM isoforms.

Downsampling done with 100K instead of 250K reads.

### Mitochondrial genes

#### Gene-level

Estimation the percentage of mitochondrial RNA captured by FLASH-seq,
using the number of reads uniquely mapped to the mitochondrial
genes.

#### Chromosome-level

Percentage of reads uniquely mapped to the mitochondrial
chromosome.

### Estimated Ribosomal Reads

Rough estimation of the percentage of ribosomal reads, using the
reads uniquely mapping to the ribosomal genes.

As this estimation does not account for multi-mapped reads, the
percentage of captured ribosomal RNA is highly under-estimated but
provides a trend that we find useful for method development
purposes.

### Gene-body Coverage

Estimated using ReSQC.

### GC Content

#### Read-level

Average read GC content per group, measured with ReSQC (directly on
FASTQ).

#### Gene-level

Average read GC content per group. Estimation based on the average GC
content of the genes from the diversity analysis.

### Gene Expression

A first overview of the relationship between the cell’s
transcriptomes. This graph is rarely (if ever) used for defining what is
the best condition.

1. Genes supported by a total of =<10 reads among all the cells are
   discarded. “MT-|MALAT1|RPS|RPL” genes are discarded
2. Depth normalisation is performed with librarySizeFactors /
   quickCluster / computeSumFactors.
3. Counts are log-normalised (logNormCounts)
4. Top 10% highly variable genes are selected (modelGeneVar,
   getTopHVGs)
5. PCA/UMAP are run (10 PCA)

### Conclusions

The number of dropouts in our control sample makes the comparison
more difficult. However, it appears that G&T-seq-like (oligo-dT only
on beads) provides equivalent results to standard FLASH-seq.

This approach can therefore be used to purify samples containing
inhibitors.

Adding both the TSO and oligo-dT to the beads decreases the reaction
efficiency.
