## Supplementary material for "Striving towards improved full-length single-cell RNA-sequencing": FLASH-seq_optimization_tests: Maxima_titration.html

Maxima Titration - Plate 580


### Maxima Titration - Plate 580

#### Maxima Titration - Plate 580

- Introduction
- Failure
  Rate
- Raw Reads
- Mapping
  Stats
- Mapping
  Errors
- Genes
  Detected
- Isoforms Detected
- Gene
  Diversity
- C.-to-C.
  corr.
- Read
  Distribution
- Gene type
- Gene Length
  - Gene-Level
  - Isoform-Level
- Mitochondrial genes
  - Gene-level
  - Chromosome-level
- Estimated Ribosomal
  Reads
- Gene-body Coverage
- GC Content
  - Read-level
  - Gene-level
- Gene
  Expression
- Conclusions

Hahaut
V., Siwicki R., Ribeiro M., Grison A. Malysheva S. & Picelli S.

05
December 2023

### Introduction

This file contains an overview of the results of our titration tests
for Maxima RT (1 ul lysis buffer).

We had previously shown that Superscript IV (ThermoFisher) and Maxima
H- (ThermoFisher) are equivalent reverse transcriptase enzymes in the
FLASH-seq protocol. This is highly beneficial for the users, as the
associated savings are significant: Superscript IV costs 0.1015-0.0593
SFr./Eur per unit of enzyme, while Maxima H- costs 0.0555-0.0288
SFr./Eur per unit of enzyme, depending on the pack size (2K vs 40K
units). Thus, on average, Superscript IV is twice as expensive as Maxima
H-.

Here, we titrated the amount of Maxima H- enzyme, to find the lowest
amount that guarantees comparable results (or better) than the original
FLASH-seq protocol.

- **p-value < 0.05**: “\*”
- **p-value < 0.01**: “\*\*”
- **p-value < 0.001**: “\*\*\*”
- **p-value < 0.0001**: “\*\*\*\*”

Whenever possible we compared the reference group
(“**Maxima\_2U**”) to the other groups using a two-sided
Wilcoxon rank sum test. Only P-values inferior to 0.05 are
displayed.

For more information regarding the analysis please contact me:
vincent.hahaut[at]iob.ch.

**Sequencing:** NextSeq550 - 75bp

**Abbreviations:**

- **Maxima 0.5U**: Maxima RT 0.5U/uL
- **Maxima 1U**: Maxima RT 1U/uL
- **Maxima 1.5U**: Maxima RT 1.5U/uL
- **Maxima 2U**: Maxima RT 2U/uL

### Failure Rate

Percentage of low quality cells after sequencing. Defined as:

### Conclusions

Based on these results, we could reduce the amount of Maxima RT from
2U to <1.5U to increase the data quality.
