## Supplementary material for "Striving towards improved full-length single-cell RNA-sequencing": FLASH-seq_optimization_tests: Miniaturization_finetuning.html

Miniaturization - Plate 582


### Miniaturization - Plate 582

#### Miniaturization - Plate 582

- Introduction
- Failure
  Rate
- Raw Reads
- Mapping
  Stats
- Mapping
  Errors
- Genes
  Detected
- Isoforms Detected
- Gene
  Diversity
- C.-to-C.
  corr.
- Read
  Distribution
- Gene type
- Gene Length
  - Gene-Level
  - Isoform-Level
- Mitochondrial genes
  - Gene-level
  - Chromosome-level
- Estimated Ribosomal
  Reads
- Gene-body Coverage
- GC Content
  - Read-level
  - Gene-level
- Gene
  Expression
- Conclusions

Hahaut
V., Siwicki R., Ribeiro M., Grison A. Malysheva S. & Picelli S.

05
December 2023

### Introduction

This file contains an overview of the results of our tests on the
lysis buffer volume (PART 1). The original FLASH-seq protocol was
performed by sorting cells in 1 ul lysis buffer followed by the addition
of 4 ul RT-PCR mix.

In principle our goal has always been to make our protocol as
accessible as possible, even to those labs that lack liquid handling
robots and/or nanodispensers. However, the downside of this approach is
the relatively large volume of reagents required for each cell.
Therefore, we explored what the minimum volume of lysis buffer is needed
to sort single cells in 384-well plates with high efficiency (i.e. high
occupancy rate). To prevent evaporation, we used a similar approach as
in the Smart-seq3xpress protocol, layering 2 ul of mineral oil on top of
the lysis buffer. We tested 4 different volumes of lysis buffer:

- **50 nl**: a 20x volume reduction compared to the
  original FLASH-seq protocol
- **100 nl**: a 10x volume reduction
- **200 nl**: a 5x volume reduction
- **300 nl**: a 3.3x volume reduction

Although one might think that the lowest volumes are the most cost
effective, this is not always the case. We have observed a higher
dropout rate (i.e., well did not contain any usable cDNA after
pre-amplification) when the amount of lysis buffer was very low. This is
likely due to the fact that cell might have not come into contact with
the lysis buffer in the first place and got lysed when trapped in the
oil phase. Although variability in droplet size between sorters exist
(FACSAria equipped with a 100 um nozzle, our standard choice, generates
4 nl droplets, while the F.SIGHT Omics generates 200 pl droplets), these
are negligible compared to the volume of lysis buffer tested here and
won’t therefore interfere with the following RT-PCR reaction.

References https://www.nature.com/articles/s41587-022-01311-4

2 μl Mineral Oil (Sigma, M5904) were first dispensed with our liquid
handling robot (Fluent, Tecan). Plates were spin down (1000g, 1 min) and
the lysis buffer dispensed on top using a nanodispenser (I-DOT,
Dispendix). Plates were spun down again before and after FACS
sorting.

- **p-value < 0.05**: “\*”
- **p-value < 0.01**: “\*\*”
- **p-value < 0.001**: “\*\*\*”
- **p-value < 0.0001**: “\*\*\*\*”

Whenever possible we compared the reference group
(“**V-300nl**”) to the other groups using a two-sided
Wilcoxon rank sum test. Only P-values inferior to 0.05 are
displayed.

For more information regarding the analysis please contact me:
vincent.hahaut[at]iob.ch.

**Sequencing:** NextSeq550 - 75bp

**Abbreviations:**

- **V-50nl**: 50 nl lysis
- **V-100nl**: 100 nl lysis
- **V-200nl**: 200 nl lysis
- **V-300nl**: 300 nl lysis

### Conclusions

The 200 nl and 300 nl conditions work well.

50 and 100 nl can be used but suffer from an increase in drop-outs
and a decrease in sensitivity. It is most likely related to the cell not
hitting the lysis buffer properly after sorting.
