## Supplementary material for "Striving towards improved full-length single-cell RNA-sequencing": FLASH-seq_optimization_tests: Nuclei_lysis_buffers.html

Polymerases - Plates 449, 478 & 480


### Polymerases - Plates 449, 478 & 480

#### Polymerases - Plates 449, 478 & 480

- Introduction
- Failure
  Rate
- Raw Reads
- Mapping
  Stats
- Mapping
  Errors
- Genes
  Detected
- Isoforms Detected
- Gene
  Diversity
- C.-to-C.
  corr.
- Read
  Distribution
- Gene type
- Gene Length
  - Gene-Level
  - Isoform-Level
- Mitochondrial genes
  - Gene-level
  - Chromosome-level
- Estimated Ribosomal
  Reads
- Gene-body Coverage
- GC Content
  - Read-level
  - Gene-level
- Gene
  Expression
- Conclusions

Hahaut
V., Siwicki R., Ribeiro M., Grison A. Malysheva S. & Picelli S.

05
December 2023

### Introduction

Here several lysis buffers were tested, with the aim of identifying
reagents capable of disrupting the nuclear membrane, thus rendering the
pre-mRNA available to the RT enzyme. Historically, RNA and DNA
extraction in bulk typically have included non-ionic detergents (e.g.,
Triton X-100, Tween-20), chaotropes (e.g., guanidium salts) or proteases
(e.g., proteinase K). High concentrations of Tween-20 or Triton X-100
(< 10% final concentration) have no influence on amplification
efficiency, whereas minute quantities of ionic detergents (0.01%) such
as sodium dodecyl sulfate (SDS) are sufficient to inhibit DNA
polymerases.

Goldenberger et al. discovered many years ago that adding at least 2%
of Tween-20 could neutralize up to 0.05% SDS and performed a very useful
titration series, which we used as a basis for our tests. In our
experiments we quenched SDS by adding 10 times excess (or higher) of
Tween-20. We also tested a range of SDS concentrations, to increase
lysis efficiency. SDS lysis was compared to the standard FLASH-seq lysis
buffer containing 0.2% Triton X-100 as well as other lysing agents
tested in previous experiments, such as guanidine hydrochloride,
sulfobetaine and DDM. We also combined NP-40 (another non-ionic
detergent), Triton X-100 and sulfobetaine, to see whether the benefits
of the different lysis buffers had an additive effect, as recently
described by Reimegård and colleagues (check also: “NO RNA DENATURATION
– PLATES 428-429”).

Reference https://genome.cshlp.org/content/4/6/368.long

- **p-value < 0.05**: “\*”
- **p-value < 0.01**: “\*\*”
- **p-value < 0.001**: “\*\*\*”
- **p-value < 0.0001**: “\*\*\*\*”

Whenever possible we compared the reference group (“**trit.
0.2%**”) to the other groups using a two-sided Wilcoxon rank sum
test. Only P-values inferior to 0.05 are displayed.

For more information regarding the analysis please contact me:
vincent.hahaut[at]iob.ch.

**Sequencing:** NextSeq550 - 75bp

**Abbreviations:**

- **Nuclei% SDS**: Nuclei, SDS 0.02%, tween
  neutralized
- **Nuclei% SDS**: Nuclei, SDS 0.01%, tween
  neutralized
- **Nuclei% SDS**: Nuclei, SDS 0.005%, tween
  neutralized
- **Nuclei% SDS**: Nuclei, SDS 0.05%, tween
  neutralized
- **SDS not neutralized**: Nuclei, SDS 0.02%, not
  neutralized
- **GuHCl**: 250 mM Guanidine Hydroxychloride
- **trit. 0.2%**: Triton 0.2%
- **NTS**: NP-40, 0.2% triton, 0.2% sulfobetaine
- **sulfo. 0.2%**: Sulfobetaine 0.2%
- **DDM**: DDM

### Failure Rate

Percentage of low quality cells after sequencing. Defined as:

- less than 100K raw reads
- more than 25% unmapped reads
- less than 75K uniquely mapped reads

This only represents the failure rate among cells that were chosen
for sequencing.

It should be noted that a large number of conditions did not work at
all.

### Raw Reads

Number of raw reads per cell.

### Mapping Stats

From STAR mapping statistics.

### Mapping Errors

From STAR mapping statistics.

### Genes Detected

### Conclusions

We observed a decrease in ribosomal / mitochondrial reads as well as
an increase in intronic reads in the SDS+tween and GuHCl conditions
suggesting that the nuclei lysis worked. The ideal condition seems to be
0.01 to 0.02% SDS + tween neutralization.
