## Supplementary material for "Striving towards improved full-length single-cell RNA-sequencing": FLASH-seq_optimization_tests: PCR_extension_time_finetuning.html

PCR duration - Plate 694 to 699


### PCR duration - Plate 694 to 699

#### PCR duration - Plate 694 to 699

- Introduction
- Failure
  Rate
- Raw Reads
- Mapping
  Stats
- Mapping
  Errors
- Genes
  Detected
- Isoforms Detected
- Gene
  Diversity
- C.-to-C.
  corr.
- Read
  Distribution
- Gene type
- Gene Length
  - Gene-Level
  - Isoform-Level
- Mitochondrial genes
  - Gene-level
  - Chromosome-level
- Estimated Ribosomal
  Reads
- Gene-body Coverage
- GC Content
  - Read-level
  - Gene-level
- Gene
  Expression
- Conclusions

Hahaut
V., Siwicki R., Ribeiro M., Grison A. Malysheva S. & Picelli S.

05
December 2023

### Introduction

In the original FLASH-seq paper we used the same pre-amplification
protocol as in Smart-seq2, with a 6-minutes long extension step. Here we
tested the optimal extension time (as others already did), in an attempt
to further reduce the sample processing time. Besides shorter
extensions, we also tried with a much longer one (10 min), to see
whether the extra time enables the KAPA DNA Polymerase to amplify even
the longest cDNA molecules.

- **p-value < 0.05**: “\*”
- **p-value < 0.01**: “\*\*”
- **p-value < 0.001**: “\*\*\*”
- **p-value < 0.0001**: “\*\*\*\*”

Whenever possible we compared the reference group (“**5 min
PCR**”) to the other groups using a two-sided Wilcoxon rank sum
test. Only P-values inferior to 0.05 are displayed.

For more information regarding the analysis please contact me:
vincent.hahaut[at]iob.ch.

**Sequencing:** NextSeq500 - 75bp

**Abbreviations:**

- **3minPCR**: 3 minutes elongation during during
  RT-PCR
- **4minPCR**: 4 minutes elongation during during
  RT-PCR
- **5minPCR**: 5 minutes elongation during during
  RT-PCR
- **6minPCR**: 6 minutes elongation during during RT-PCR
  (Plate 580)
- **10minPCR**: 10 minutes elongation during during
  RT-PCR

Of note, our 6 min PCR plate from this batch did not work as expected
and was replaced by plate 580 from another batch. Small variations can
be therefore expected.

### Failure Rate

Percentage of low quality cells after sequencing. Defined as:

### Conclusions

We observe that 10 minutes provides the best results overall,
suggesting that increaseing the PCR duration may still have some
benefits. However, these improvements are rather minor compared to stark
increase in processing time. We therefore recommend using 4-5 minutes
PCR which provides a better balance.
