## Supplementary material for "Striving towards improved full-length single-cell RNA-sequencing": FLASH-seq_optimization_tests: Polymerases_comparison.html

Polymerases - Plate 481 & 508


### Polymerases - Plate 481 & 508

#### Polymerases - Plate 481 & 508

- Introduction
- Failure
  Rate
- Raw Reads
- Mapping
  Stats
- Mapping
  Errors
- Genes
  Detected
- Isoforms Detected
- Gene
  Diversity
- C.-to-C.
  corr.
- Read
  Distribution
- Gene type
- Gene Length
  - Gene-Level
  - Isoform-Level
- Mitochondrial genes
  - Gene-level
  - Chromosome-level
- Estimated Ribosomal
  Reads
- Gene-body Coverage
- GC Content
  - Read-level
  - Gene-level
- Gene
  Expression
- Conclusions

Hahaut
V., Siwicki R., Ribeiro M., Grison A. Malysheva S. & Picelli S.

05
December 2023

### Introduction

KAPA HiFi HotStart ReadyMix has been shown already 12 years ago by M.
Quail and collaborators to be among the best enzymes for unbiased DNA
amplification. In an attempt to find an even better one for our
applications, we tested several new DNA Polymerases not available when
that study was published. It must be kept in mind that in FLASH-seq we
combine RT and PCR, therefore not only does the DNA Polymerase need to
show a superior performance to KAPA HiFi but also the reverse
transcriptase needs to work in a different PCR buffer. This reduced the
number of viable candidates considerably. Only the enzymes that worked
at least in one buffer combination are shown here.

References https://www.nature.com/articles/nmeth.1814

- **p-value < 0.05**: “\*”
- **p-value < 0.01**: “\*\*”
- **p-value < 0.001**: “\*\*\*”
- **p-value < 0.0001**: “\*\*\*\*”

Whenever possible we compared the reference group (“**Kapa
HS**”) to the other groups using a two-sided Wilcoxon rank sum
test. Only P-values inferior to 0.05 are displayed.

For more information regarding the analysis please contact me:
vincent.hahaut[at]iob.ch.

**Sequencing:** NextSeq550 - 75bp

**Abbreviations:**

- **Kapa HS**: Kapa Hifi 2x Master Mix (Roche)
- **seqAmp**: seqAmp DNA polymerase (Takara)
- **Phusion**: Phusion Hot Start II DNA Polymerase (2
  U/µL) (Thermofisher)
- **Equinox**: Equinox library amplification kit
  (Watchmaker)
- **SuperFi**: Platinum™ SuperFi™ DNA Polymerase
  (Thermofisher)
- **SuperFi 2x**: Platinum™ SuperFi™ PCR Master Mix
  (Thermofisher)
- **Kapa2G+GC**: Kapa2G + high GC buffer (Roche)
- **Q5**: Q5® High-Fidelity 2X Master Mix (NEB)
- **Kapa2G+B**: Kapa2G + high B buffer (Roche)
- **Kapa2G+A**: Kapa2G + high A buffer (Roche)
- **Kapa2G+MM**: Kapa2G + high MM buffer (Roche)

### Conclusions

As previously observed, Kapa Hifi 2x master mix provides the best
results. However Equinox (Watchmaker) comes in close second.

The other polymerase did not reach sufficient data quality.
