## Supplementary material for "Striving towards improved full-length single-cell RNA-sequencing": FLASH-seq_optimization_tests: ProteinaseK_for_lysis.html

ProteinaseK - Plate 609


### ProteinaseK - Plate 609

#### ProteinaseK - Plate 609

- Introduction
- Failure
  Rate
- Raw Reads
- Mapping
  Stats
- Mapping
  Errors
- Genes
  Detected
- Isoforms Detected
- Gene
  Diversity
- C.-to-C.
  corr.
- Read
  Distribution
- Gene type
- Gene Length
  - Gene-Level
  - Isoform-Level
- Mitochondrial genes
  - Gene-level
  - Chromosome-level
- Estimated Ribosomal
  Reads
- Gene-body Coverage
- GC Content
  - Read-level
  - Gene-level
- Gene
  Expression
- Conclusions

Hahaut
V., Siwicki R., Ribeiro M., Grison A. Malysheva S. & Picelli S.

05
December 2023

### Introduction

To improve gene capture, we modified the lysis procedure to include a
mild protease digestion, similar to what recently described by
Rodriguez-Meira and collaborators. In our tests we used 0.25 uL of
thermolabile proteinase K (NEB, cat. # P8111S), which we added at a
concentration of 0.006 U/ul in the lysis buffer. The enzyme was
subsequently heat inactivated to avoid inhibition of the RT and PCR
enzymes (10 min at 72 °C).

References https://www.sciencedirect.com/science/article/pii/S1097276519300097

- **p-value < 0.05**: “\*”
- **p-value < 0.01**: “\*\*”
- **p-value < 0.001**: “\*\*\*”
- **p-value < 0.0001**: “\*\*\*\*”

Whenever possible we compared the reference group
(“**FS-Std**”) to the other groups using a two-sided
Wilcoxon rank sum test. Only P-values inferior to 0.05 are
displayed.

For more information regarding the analysis please contact me:
vincent.hahaut[at]iob.ch.

**Sequencing:** NextSeq550 - 75bp

**Abbreviations:**

- **FS-Std**: FLASH-seq.
- **ProteinaseK**: FLASH-seq with Proteinase K in
  lysis.

### Failure Rate

### Conclusions

The addition of thermolabile proteinase K did not notably impact the
reaction. One follow up experiment could be to test the gDNA
accessibility as described in TARGET-seq (https://www.sciencedirect.com/science/article/pii/S266616672030112X?via%3Dihub).
