## Supplementary material for "Striving towards improved full-length single-cell RNA-sequencing": FLASH-seq_optimization_tests: RNA_denaturation_conditions_various_LB.html

Lysis Buffer without denaturation - Plates 427/428


### Lysis Buffer without denaturation - Plates 427/428

#### Lysis Buffer without denaturation - Plates 427/428

- Introduction
- Raw Reads
- Mapping
  Stats
- Mapping
  Errors
- Genes
  Detected
- Isoforms Detected
- Gene
  Diversity
- C.-to-C.
  corr.
- Read
  Distribution
- Gene type
- Gene Length
  - Gene-Level
  - Isoform-Level
- Mitochondrial genes
  - Gene-level
  - Chromosome-level
- Estimated Ribosomal
  Reads
- Gene-body Coverage
- GC Content
  - Read-level
  - Gene-level
- Gene
  Expression
- Conclusions

Hahaut
V., Siwicki R., Ribeiro M., Grison A. Malysheva S. & Picelli S.

05
December 2023

### Introduction

Reimegård and collaborators recently showed that, by changing the
lysis buffer composition and removing the heat denaturation step prior
RT reaction, it is possible to simultaneously measure gene expression
and intracellular proteins in individual cells. In that paper they used
a modified Smart-seq2 protocol for scRNA-seq. Due to the similarity
between the Smart-seq2 and FLASH-seq mixes, we thought that such
modified buffer could also be beneficial in our settings. Besides the
commonly used FLASH-seq lysis buffer ( 0.2% Triton X-100 and 50 mM
guanidine hydrochloride), we also tested 0.2% sulfobetaine
(3-(1-Pyridinio)-1-propanesulfonate, Sigma cat. # 82804-50G) and a
combination of 3 different detergents (NTS: 1% NP-40, 0.1% Triton X-100,
0.1% sulfobetaine), also described by Reimegård and collaborators. For
completeness, the 4 lysis buffers were tested with and without heat
denaturation.

References https://www.nature.com/articles/s42003-021-02142-w

- **p-value < 0.05**: “\*”
- **p-value < 0.01**: “\*\*”
- **p-value < 0.001**: “\*\*\*”
- **p-value < 0.0001**: “\*\*\*\*”

Whenever possible we compared the reference group (“**72°C
3min, trit. 0.2%**”) to the other groups using a two-sided
Wilcoxon rank sum test. Only P-values inferior to 0.05 are
displayed.

For more information regarding the analysis please contact me:
vincent.hahaut[at]iob.ch.

**Sequencing:** NextSeq550 - 75bp

**Abbreviations:**

With 3 minutes 72°C denaturation.

- **no denat. GuHCl**: Guanidine Thiocyanate at 250 mM in
  lysis.
- **no denat. trit. 0.2%**: Triton X-100 0.2%.
- **no denat. sulfo. 0.2%**: Sulfobetaine 0.2%.
- **no denat. NTS**: 1% NP-40, 0.1% Triton X-100, 0.1%
  sulfobetaine (NTS).

Without denaturation.

- **72°C 3min GuHCl**: Guanidine Thiocyanate at 250 mM in
  lysis.
- **72°C 3min trit. 0.2%**: Triton X-100 0.2%.
- **72°C 3min sulfo. 0.2%**: Sulfobetaine 0.2%.
- **72°C 3min NTS**: 1% NP-40, 0.1% Triton X-100, 0.1%
  sulfobetaine (NTS).

### Conclusions

10x Genomics lysis buffer does not require denaturation. We tested
different lysis buffers to see if denaturation could be skipped.

In general, no denaturation resulted in a greater number of failed
cells. Moreover, the data quality decrease as well compared to our
regular 72°C 3 min with 0.2% Triton X-100.

Interestingly the number of detected genes did not decrease when
skipping denaturation. However, we also observe more intronic/intergenic
reads.

Moreover, the data quality decreased as well compared to our regular
72°C 3 min with 0.2% Triton X-100. However, skipping the denaturation
can also be done with minimal drawbacks.
