## Supplementary material for "Striving towards improved full-length single-cell RNA-sequencing": FLASH-seq_optimization_tests: TCEP_to_replace_DTT.html

TCEP Titration - Plate 421


### TCEP Titration - Plate 421

#### TCEP Titration - Plate 421

- Introduction
- Failure
  Rate
- Raw Reads
- Mapping
  Stats
- Mapping
  Errors
- Genes
  Detected
- Isoforms Detected
- Gene
  Diversity
- C.-to-C.
  corr.
- Read
  Distribution
- Gene type
- Gene Length
  - Gene-Level
  - Isoform-Level
- Mitochondrial genes
  - Gene-level
  - Chromosome-level
- Estimated Ribosomal
  Reads
- Gene-body Coverage
  - Gene-level
- Gene
  Expression
- Conclusions

Hahaut
V., Siwicki R., Ribeiro M., Grison A. Malysheva S. & Picelli S.

05
December 2023

### Introduction

This file contains an overview of the results of our titration of
TCEP to replace DTT (1 ul lysis buffer).

In the patent WO2014210353A2 the Authors tested different reducing
agents for dissolution of hydrogel beads in the context of droplet
RNA-seq (10x Genomics). TCEP (tris(2-carboxyethyl)phosphine, Thermo
Fisher cat. # 77720) was found to be as effective as DTT but, unlike
DTT, it is odorless, very soluble in aqueous buffers at nearly any pH
and very stable. It was also reported to require lower concentrations
compared to DTT. For comparison, DTT is used in FLASH-seq at a 5 mM
final concentration in the RT-PCR reaction but here we explored a larger
interval of lower and higher concentrations.

- **p-value < 0.05**: “\*”
- **p-value < 0.01**: “\*\*”
- **p-value < 0.001**: “\*\*\*”
- **p-value < 0.0001**: “\*\*\*\*”

Whenever possible we compared the reference group (“**trit.
0.2%**”) to the other groups using a two-sided Wilcoxon rank sum
test. Only P-values inferior to 0.05 are displayed.

For more information regarding the analysis please contact me:
vincent.hahaut[at]iob.ch.

**Sequencing:** NextSeq550 - 75bp

**Abbreviations:**

Except if stated otherwise, the lysis buffer contained sulfobetaine
0.2% instead of Triton X-100 0.2%.

- **TCEP 1mM**: TCEP 1 mM.
- **TCEP 2.5mM**: TCEP 2.5 mM.
- **TCEP 5mM**: TCEP 5 mM.
- **TCEP 10mM**: TCEP 10 mM.
- **trit. 0.2%**: Regular FLASH-seq.
- **sulfo. 0.2%**: Regular FLASH-seq with sulfobetaine
  lysis buffer instead of triton X-100.

### Failure Rate

### Conclusions

It seems that TCEP (1 mM) can be used as a less toxic substitute to
replace DTT.

However, it also resulted in a significantly lower number of cells
passing filters (30%).

These results will require a second test with a different plate to
ensure that it is not due to a sorting issue.
